# A Ubiquitin network safeguards cell identity by continuously degrading a stem-cell related translational machinery

**DOI:** 10.64898/2026.03.06.709266

**Authors:** Salwa Danial, Raya Ghanem, Muhammad Makhzumy, Eliya Bitman-Lotan, Jelly Soffers, Jesslyn C. Henriksen, Avital Sarusi-Portuguez, Olivia S. Rissland, Ryan D. Mohan, Ayala Shiber, Amir Orian

## Abstract

How cell identity is maintained is a fundamental question, and loss of cell identity is a hallmark of aging that is associated with multiple age-related diseases. In the adult *Drosophila* midgut, we identified a post-transcriptional regulatory layer that supervises enterocyte (EC) identity and fails upon aging. Combining single-cell RNA-seq with lineage tracing in aging ECs and classical genetics we found that aging ECs express genes that unlock the differentiated state. Upon aging, the protein Rogue (CG13928), a translational repressor, orchestrates reactivation of a stem-cell-related translational repression machinery involving p-body-associated RNA-binding proteins that cancels the differentiated state. In young ECs, this machinery is continuously suppressed by the deubiquitinase Non-stop (dUSP22) and the ubiquitin E2 dUbcH8/Kdo and the E3 enzyme CTLH, together suppress the stem-cell related RNA binding proteins safeguarding EC identity. Upon aging, the levels of dUSP22 decline, dUbcH8/Kdo and the E3 CTLH complex are cleared via Rogue, and the stem cell-related p-bodies are reactivated, self-destroying EC identity.

## Introduction

### Continuous supervision of cell identity

How cell identity is maintained is a fundamental question. As postulated long ago, regulation of differentiated cell identity requires continuous supervision by “identity supervisors”(ISs)^1^. ISs enhance expression of genes required for executing the physiological tasks of the differentiated cell while suppressing expression of previous and non-relevant gene programs^2–7^. Inability to safeguard the differentiated state is a hallmark of aging, and results in diseases such as neurodegeneration, metabolic disorders, and cancer^8–12^. Moreover, reprogramming differentiated cells to induced pluripotent stem cells (iPS) is inefficient due to intrinsic barriers within differentiated cells that maintain identity and prevent de-differentiation or trans-differentiations^13–17^. Loss of identity is observed upon aging due to both a decline in identity supervisor levels and pathological expression of identity breaker (IB) genes. IBs unlock cell identity and accumulate pathologically upon aging ^16,17^, and unlocking cell identity is also a fundamental step in tumorigenesis^11,12^. Well-known examples of IBs are MyoD which converts fibroblasts to muscle progenitors^18,19^, and Yamanaka factors (OSKM: Oct3/4, Sox2, Klf4, c-Myc) which convert fibroblasts into induced pluripotent cells (iPS) ^20,21^.

### *Drosophila* midgut, and enterocyte identity

The intestinal epithelium is a powerful model for studying the regulation of cell identity, as well as aging. In both mammals and *Drosophila* the adult gut is a highly regenerating tissue^22–25^. In the fly midgut, intestinal stem cells (ISCs) proliferate to self-renew or give rise to mature differentiated gut enterocytes (ECs) and enteroendocrine (EE) cells (Fig. 1A) ^25–28^. Cell identity is regulated in part at the chromatin and transcriptional levels.^3,17,29,30^. In differentiated ECs, the transcription factor Hey together with type A lamin, Lamin-C (LamC), or the nuclear non-stop identity complex (NIC) safeguard EC identity by preventing expression of progenitor and non-relevant programs ^16, 17^. Eliminating identity supervisors (such as Hey or subunits of NIC) or expressing identity breakers in young ECs (such as the ISC-related LamDm0) disrupts cell identity and reduces overall survival. Likewise, expression of identity supervisors, or elimination of identity breakers in ECs of aging animals suppresses aging and extends longevity^16,17^. However, the post-transcriptional mechanisms that maintain EC identity and silence identity breakers in differentiated cells are largely unknown. It is also unclear how these mechanisms are perturbed and deregulated upon aging.

**Figure 1:**
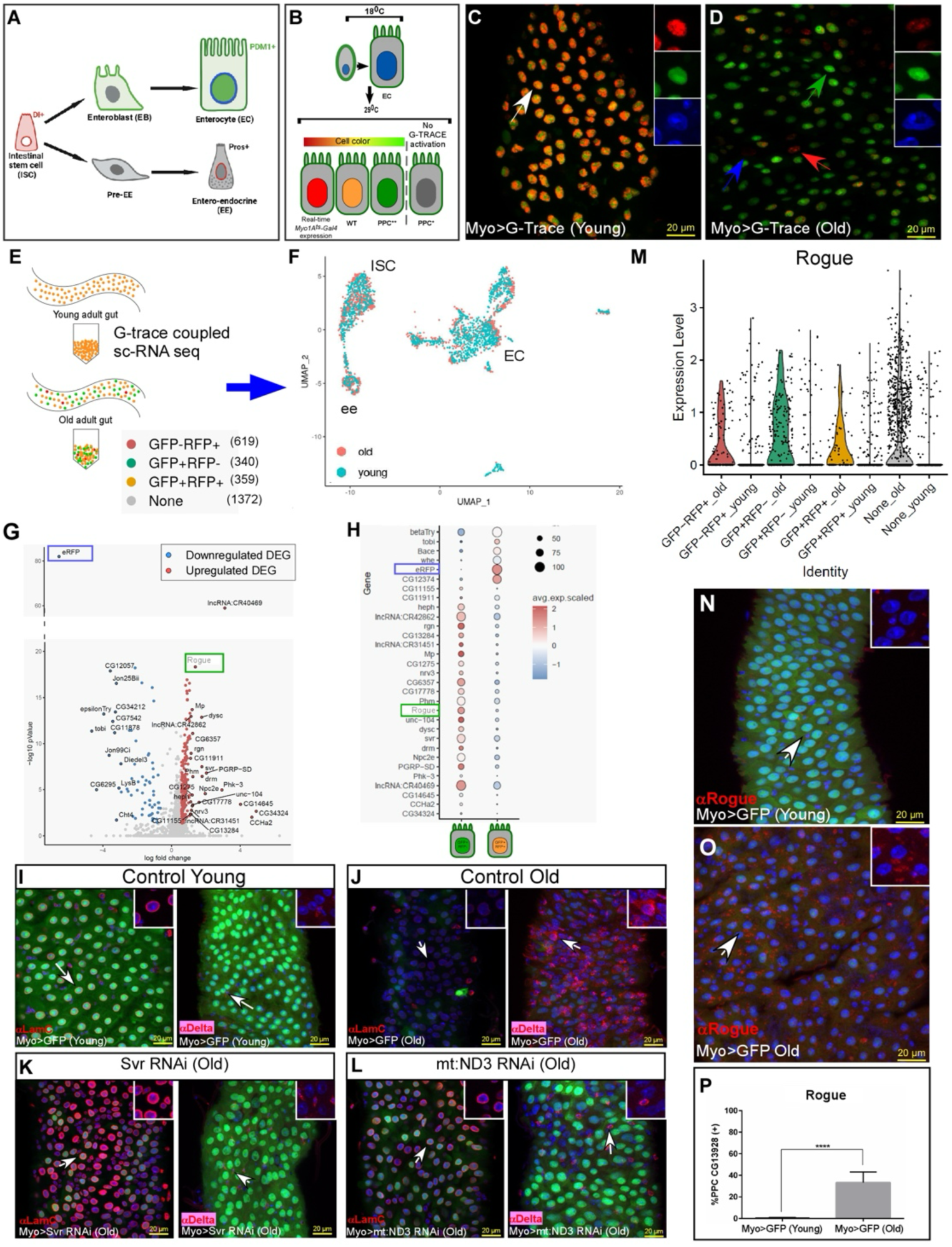
sc-RNA sequencing coupled with G-TRACE linage tracing identifies aged-related potential IBs in ECs. **(A)** Cartoon depicting the cells that populate the adult *Drosophila* midgut. Abbreviations: Dl: Delta, Pdm1: Nubbin, Pros: Prospero, ISC: intestinal stem cells, EC: Enterocyte, EE: enteroendocrine cell. **(B)** Diagram of the MyoIA>Gal4/Gal80^ts^∼G-TRACE linage tracing in the *Drosophila* midgut (see text for details). PPC* are likely miss-differentiated progenitors where the G-TRACE system was not activated (gray). **(C, D)** Confocal images of young and aged ECs expressing G-TRACE system. (C) In young midguts all ECs express both RFP and GFP and are, therefore, orange. (D) Five weeks old guts (old) exhibit diverse cell populations including GFP+RFP^-^ cells (PPC** are EC that are no longer differentiated). In both C and D, arrows point to cells shown in insets, and DAPI (Blue) marks DNA. Scale bar is 20μM. **(E)** Experimental design of the G-TRACE∼scRNA-Seq experiment in old and young ECs. **(F)** Classification of single cells into distinct cell populations in young and aged midguts according to gene expression (32). **(G, H)** Expression of transcripts in PPC** ^GFP+ RFP-^ in old midguts, compared to young EC^GFP+ RFP+^. (G) Volcano plot of significantly upregulated (•) and downregulated (•) DEGs comparing young ECs to PPC**. (H) Examples of genes exhibiting increased (red) or reduced (blue) expression in PPC**^GFP+, RFP-^ compared to EC^GFP+, RFP+^. The RFP is donated in a blue box. A green box highlights the translational repressor Rogue (CG13928). **(I-L)** RNAi-mediated elimination of the indicated genes exhibiting ectopic mRNA upregulation in aged PPC** suppresses aging phenotypes of aged EC-like cells. Confocal images of young midguts (I), or five weeks old-aged midgut tissue expressing control (J), or ECs conditionally expressing the indicated UAS transgenes using the MyoIA>Gal4/Gal80^ts^ system (K, L). MyoIA>UAS-GFP marks fully differentiated ECs, LamC or Delta are shown in red, and DAPI (blue) marks DNA. White arrows point to cells shown in insets. Scale bar is 20μM. **(M)** Violin plot of *Rogue* (CG13928) mRNA distribution in the different aged midguts polyploid cells expressing G-TRACE, where each dot represents a single cell. **(N-P)** Rogue protein (red), is not expressed in young ECs (N), but is ectopically expressed in aged ECs (O). (P) Quantification of Rogue (CG13928) protein abundance in young and old ECs, ****=p<0.0001.

## Results

We hypothesized that the aging gut contains EC-like cells, that are no longer differentiated, which contribute to loss of tissue homeostasis, and that these cells arise through ectopic expression of identity breakers. To identify and visualize these no-longer-differentiated ECs, we used G-TRACE lineage tracing^17,31^ (Fig. 1B). G-TRACE combines the enterocyte-specific Myo1A-Gal4/Gal80 with a transgene expressing UAS-RFP, UAS-Flpase, and a Flp-out eGFP tracer. The Myo1A-Gal4 is active in parental differentiated ECs and activates a UAS-RFP that indicates Gal4 activity in real time. Concomitantly, in the parental EC, the Gal4 generates a “Flp-out” event, resulting in activation of the eGFP tracer from a ubiquitous promoter (Ubi63FRT>stop>FRTeGFP) regardless of the current cell identity. In young guts, all ECs express both RFP and GFP (Fig.1C). However, in aging guts, multiple types of polyploid EC-like cells are observed, including misdifferentiated progenitors that did not activate the tracing system (DAPI only positive depicted in gray in 1B). Of importance are “GFP-only” expressing cells that are likely ECs which are no-longer differentiated^16,17^ (termed PPC**; Figure 1D green arrow).

To identify the gene signature of these PPC** and the potential IBs they express, we combined single-cell RNA-seq (sc-RNAseq) with G-TRACE analysis. We used young guts as a control for profiling normal EC cell populations (GFP^+^, RFP^+^) and observed that many of these populations correspond well with published scRNA-seq data and the midgut cell atlas (Fig. 1E, F)^32^. sc-RNA-seq analysis of these PPC** revealed reduced expression of genes involved in the physiological functions of ECs such as proteolysis, serine peptidases, and sugar metabolism (Figs. 1G, S1A, B, Table S1). Remarkably these PPC** exhibited 205 enriched transcripts belonging to pathways related to prior, or other cell fates such as midgut progenitors (*escargot, toy*, *sox21A, dmyc*), neurogenesis and signaling (Figs. 1G, H, S1C, examples are shown in Fig. S1, and the complete list of differentially expressed genes (DEG) under Table S1). We hypothesized that some of these upregulated genes, observed solely in PPC**, may be potential IB genes. Therefore, we tested whether eliminating potential IB genes was able to suppress loss of identity phenotypes observed in aged EC. Aged ECs are characterized by reduced expression of LamC, ectopic expression of the ISC marker and Notch ligand, Delta and exhibit loss of tissue homeostasis and gut integrity^17,33,34^. Indeed, RNAi-mediated elimination of seven out of nine of these potential IB genes from aged ECs, restored LamC expression and suppressed the ectopic expression of Delta (Figure 1I-L, Supp. Fig.2A-K).

**Figure 2:**
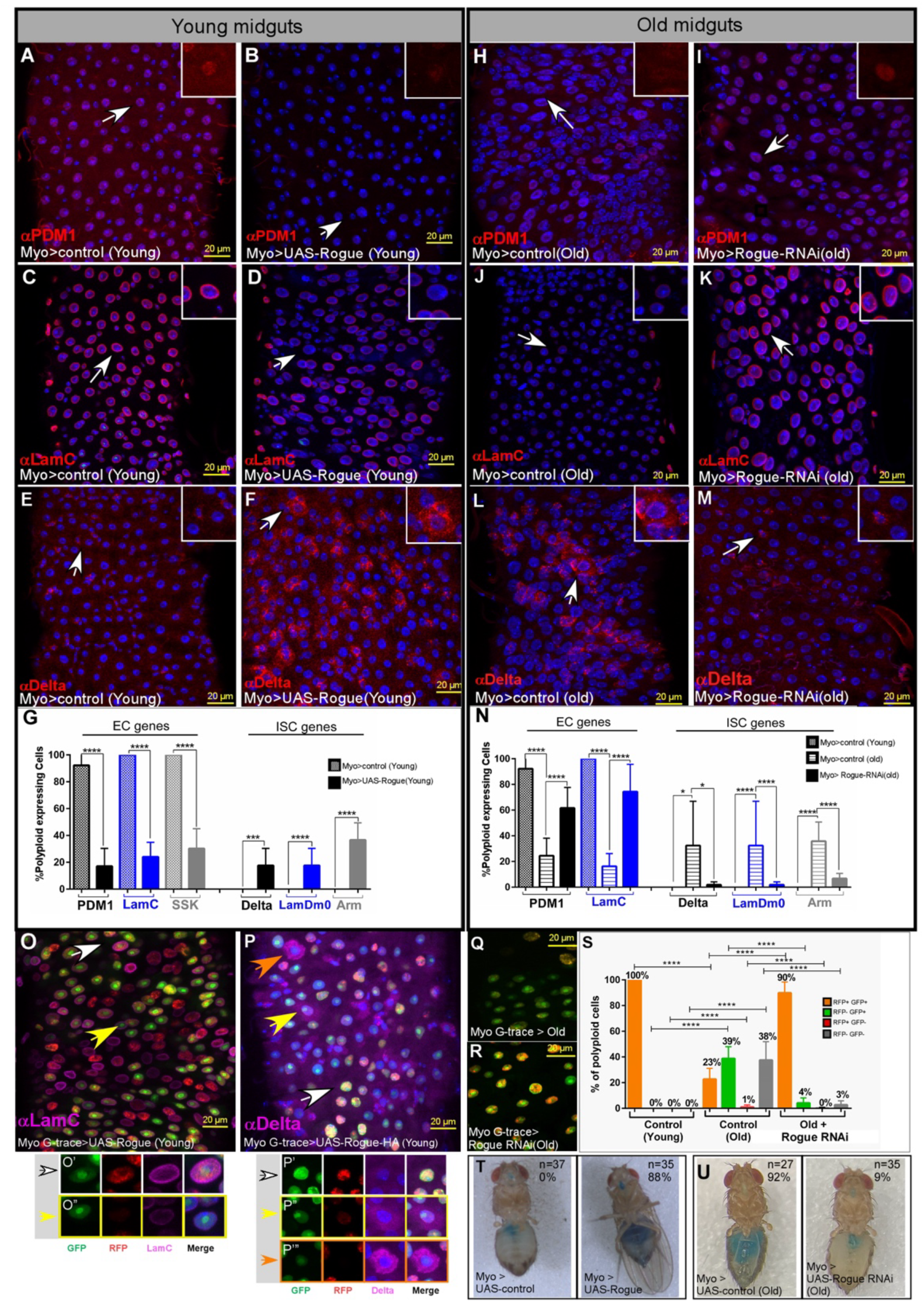
Rogue (CG13928) is an EC identity breaker (IB). **(A-G)** Expression of Rogue (CG13928) in young ECs cancels EC identity. Confocal images of young midguts expressing control, or Rogue in ECs using the MyoIA>GAL4/Gal80^ts^ system and the indicated antibodies. Arrows point to cells shown in insets and DAPI (blue) marks DNA. (A, B) PDM1; (C, D) LamC; (E, F) Delta; (G) Quantification of 3 independent biological repeats, ****=p<0.0001 ***=p<0.001. **(H-N**) Elimination of Rogue from aged ECs suppresses aging phenotypes and restores EC identity. Confocal images and quantification of five weeks old ECs expressing control (H, J, L), or UAS-Rogue RNAi (I, K, M) using the MyoIA>GAL4/Gal80^ts^ system and the indicated antibodies. Arrows point to cells shown in insets and DAPI (blue) marks DNA, Scale Bar is 20μM. (N) Quantification of 3 biological repeats ****=p<0.0001 *=p<0.1. (**O-S)** G-TRACE analysis of the effects on EC identity of Rogue over-expression in young ECs or its elimination from aged ECs. (O-P) Confocal images of ECs expressing UAS-Flag-Rogue (CG13928) in enterocytes using the MyoIA>GAL4/Gal80^ts^-GTRACE coupled system and the indicated antibodies. (O) LamC, (P) Delta. DAPI (Blue) marks DNA, White arrow points to EC^GFP+RFP+^ cells, yellow and orange arrows point to PPC**^GFP+RFP-^. Scale bar is 20μM, and magnifications are shown below (O’, O”, P’, P”). **(Q-S)** G-TRACE analysis of aged ECs, or aged EC where Rogue (CG13928) was eliminated using UAS-RNAi. (S) Quantification of the distribution of polyploid cells using G-TRACE of young ECs, or aged ECs, or aged EC where Rogue was eliminated using RNAi. **(T, U)** Expression of UAS-Rogue in young ECs (T, right panel) but not control (T, left panel) impairs gut integrity as evident by Smurf assay. (U) Rogue elimination from aged EC using UAS-Rogue RNAi restores gut integrity as evident by “Smurf” assay ****=p<0.0001.

### Rogue (CG13928) is a bona-fide identity breaker

One gene of interest was CG13928, that we termed *Rogue,* a translational repressor that was not present in young ECs, but its mRNA and protein levels accumulated in aged PPC** (Figs. 1G, H, 1M-P Table S1). Rogue (CG13928) protein was also not expressed in young or aged differentiated enteroendocrine cells (Ees, Fig S2L, M). Therefore, we focused our study on the role of Rogue in aged ECs. Rogue was originally identified as a translational repressor interacting with monomeric translational repressor Orb2 to co-repress translation in the *Drosophila* brain^35^. In the midgut, *rogue* mRNA was ectopically upregulated in young ECs upon eliminating non-stop a deubiquitinase (DUB) that is part of the identity supervising nuclear complex termed NIC^16^.

We tested whether Rogue is a bona-fide IB using both gain- and loss-of-function experiments (GOF, LOF, respectively). Forced expression of Rogue in young ECs canceled EC identity; it reduced the expression of the EC founder transcription factor PDM1 (Nubbin), LamC, the EC-specific cell adhesion molecule Snakeskin (SSK), and the MyoIA>GFP EC marker. Expression of Rogue was also sufficient to drive ectopic expression of ISC genes such as Delta, LamDm0, and the ISC-specific surface protein and β−catenin ortholog Armadillo (Arm) (Figs. 2A-G, S3A-G). Moreover, elimination of Rogue in five-week-old ECs suppressed aging phenotypes and restored EC identity. RNAi-mediated loss of Rogue using multiple transgenic lines restored the expression of Pdm1 and LamC (Figs. 2H-K, N, S3, S4A-D). It also suppressed age-related ectopic expression of ISC genes such as Delta, LamDm0 and Arm (Figs. 2L-N, S3H-L, S4E-H). We observed similar findings using G-TRACE analysis; expression of Rogue resulted in loss of LamC and ectopic expression of Delta in PPC** (Fig. 2O-P”). Moreover, depletion of Rogue by RNAi in aged ECs greatly restored the G-TRACE-based cell distribution of aged ECs, similar to that observed in young ECs; Loss of Rogue increased the number of GFP^+^RFP^+^ cells concomitantly with a reduction in the number of PPC** (GFP^+^RFP^-^) and mis-differentiated progenitors (GFP^-^, RFP^-^polyploid cells; Fig. 2Q-S). Similarly to Rogue, we found that its binding partner, the Orb2 protein accumulates in aged ECs, but not aged EEs, and that Orb2 is likely an identity breaker as evident in both GOF and LOF experiments (not shown). At the tissue level, expression of Rogue, but not control, in young ECs resulted in loss of gut integrity, as measured by the Smurf assay; extravagation of blue-colored food outside the gut to the entire body (Fig. 2T)^16, 17^. Likewise, elimination of Rogue in aged ECs restored gut integrity (Fig. 2U).

### Rogue disrupts EC translation

Rogue is a translational repressor, yet little is known regarding the effect(s) of Rogue on the translational machinery. As a starting point, we determined the impact of Rogue on the cellular polysome landscape upon expression of FLAG-Rogue in young ECs^36,37^. We conditionally expressed UAS-FLAG-Rogue, or UAS-GFP control and preformed polysome profiling of extracts derived from abdomen of young guts (see details under methods). Expression of FLAG-Rogue in ECs shifted translation globally, as determined by a reduction in polysomes/monosomes ratio (Figs.3 A-C, S5A). We observed a similar shift in monosome/polysome ratio in extract derived from abdomens of five weeks aged animals. This shift was suppressed in animals expressing UAS-Rogue RNAi in ECs using the MyoIA>Gal4/Gal80^ts^ system (Figs. 3D, 3E, S4I-L S5B) indicating that the translation levels of ECs are recovered in aged animals in ECs expressing RNAi targeting Rogue. Western blot analysis revealed that FLAG-Rogue is associated with ribosome subunits, predominantly the 40S-60S and monosome fractions but not in the heavy translated polysomes fractions (Fig. S6C). These observations suggest that Rogue may be involved in repression of translation.

**Figure 3:**
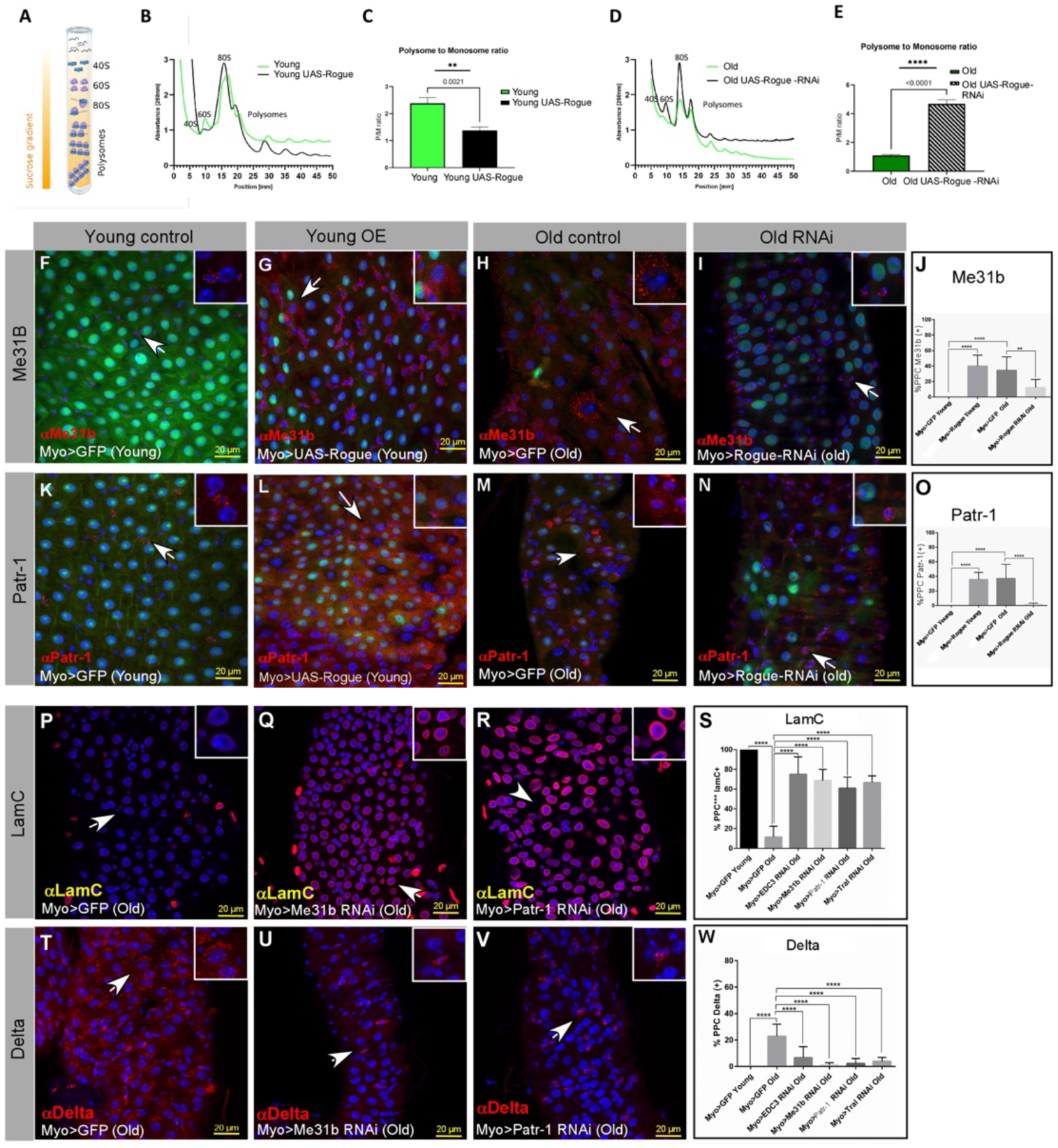
Rogue impairs the distribution of ribosome subunits, induces p-bodies proteins that ectopically accumulate in ECs upon aging and are potential IBs. **(A)** Polysome profiling; schematic view of monosomes and polysomes distribution over a sucrose gradient. **(B-E)** Representative polysome profiling of control, or young midguts expressing control UAS-Rogue (B, C), or five weeks old midguts expressing control, or UAS-Rogue RNAi in EC (D, E). (C) and (E) are quantifications of polysome/monosome ratios (P/M). In both experiments three biological repeats were performed and *** *p*<0.0004, and **= *p.= 0.00345* respectively. (F, G, K, L) p-body-proteins levels in young midguts expressing control (F, K) or UAS-Rogue (G, L), and quantification is shown in **(J, O). (H, I, M, N)** Me31B and Patr-1 proteins accumulated in aged midguts (H, M), but not in ECs of aged midguts expressing UAS-Rogue RNAi in ECs (I, N). Quantification is shown in (J, O) and in all experiments n=3 and ****=p<0.0001. **(P-W)** P-body-proteins are potential IBs. Confocal images of aged midguts where the indicated P-body-proteins were eliminated from aged ECs using the indicated UAS-RNAi and MyoIA>GAL4/Gal80^ts^ expression system. Expression of the indicated RNAi restores LamC expression (P-S) and cancels the ectopic expression of Delta (T-W).

Complementing these observations, using G-TRACE coupled scRNAseq and comparing young ECs to ECs that express Rouge we find that Rouge expression results in suppression of ECs gene signatures. Concomitantly it induces the ectopic expression of ISC, EE and other fate programs. Yet these cells do not become functional stem cells or trans-differentiate to EEs (Fig. S5D-G and see discussion).

### A stem-cell related translational repression machinery is reactivated by Rogue and upon aging

Processing bodies/p-granules are membraneless cytoplasmic organelles that serve as key sites of post-transcriptional mRNA control, functioning, in part, by harboring translation repressors and mRNA decay pathways^38–40^. In young *Drosophila* midguts, P-bodies are present in ISCs, inhibiting the expression of key differentiated EC mRNAs such as Pdm1, safeguarding ISC identity and preventing premature differentiation. Upon differentiation, P-bodies somehow dissolve, enabling translation of differentiated EC genes^41^. Interestingly, Rogue was identified in a screen as a positive regulator of P-body shape in ISCs^41^. Moreover, a shift in the profile from polysomes to monosomes is known to correlates with the emergence of P-bodies^42^. Indeed, while in young midguts the expression of P-body proteins is confined to ISCs, we found that expression of Rogue or Orb2 in young ECs but not in EE, resulted in pathological accumulation of core P-body proteins such as Me31B and Part-1 (Figs. 3F, G, K, L and quantified in Fig. 3J, O; Fig. S6K, L, and not shown). Moreover, in aging guts P-bodies are present in EC-like polyploid cells (PPC**), and elimination of Rogue or Orb2 from aged ECs suppressed the ectopic expression of P-body proteins (Figs. 3H, I, M, N; quantified in Fig.3 J, O and not shown). Likewise, we observed that the ectopic expression of P-body-proteins such as Me31B in aged ECs was suppressed by RNAi-mediated elimination of IBs identified in our G-TRACE coupled scRNA-seq (Fig. S6A-D). Remarkably, we found that core P-body proteins are potential IBs, as RNAi-mediated elimination Me31B, or Patr-1, or TRAL, or EDC3 in aged ECs restored LamC expression and suppressed the ectopic expression of ISC marker Delta (Figs.3P-W; S6E-J). Thus, aging ECs ectopically reactivate P-bodies proteins, and along with Rogue, “self-destruct” EC identity.

### Continuous silencing of Rogue and P-body proteins in young ECs is mediated by a dedicated E2/E3 ubiquitin complex

Rogue and core P-body-proteins are active in stem cells (ISCs), preventing premature differentiation^41^. They are, however, silenced by unknown mechanism(s) in young ECs and re-accumulate in aged ECs. At the protein level, elimination of the identity supervisor non-stop from young ECs results in ectopic expression of both Rogue as well as core P-body proteins (Figs.4A-E, S7A, B). Moreover, overexpression of non-stop in aged ECs suppresses ectopic expression of Rogue and core P-body-proteins, extending longevity (Fig. S7C-I). Previously, we found that *Rogue* mRNA is transcriptionally repressed in young ECs by non-stop^16^. However, unlike *Rogue*, the levels of P-body-protein mRNAs were not upregulated in aged PPC** or upon elimination of non-stop^16^ (Table S1). Taken together, these observations suggest non-stop represses *Rogue* at the transcriptional/mRNA level in young ECs, while it silences P-body proteins indirectly via a different mechanism.

One mechanism that may silence P-body-proteins in young ECs is their degradation by the ubiquitin proteasome system (UPS). Indeed, in the developing embryo, the E2 dUbCE2H also termed Marie Kondo (Kdo) and the carboxy-terminal to LisH (CTLH) E3 RING-type Ub-ligase complex degrade P-body proteins such as Me31B and TRAL during maternal to zygotic transition (MTZ) ^43,44^. The Kdo∼CTLH E3 ligase is a macromolecular multi-subunit E3 ubiquitin ligase that mostly functions as a dimer and is highly conserved in yeast (GID complex), flies, and humans (Fig. 4F) ^45^. We speculated that in young ECs, the Kdo∼CTLH complex is required to continuously eliminate P-body-proteins. To test this hypothesis, we eliminated the Kdo or Ran-BPM (a subunit of CTLH) from young ECs and observed ectopic accumulation of Rogue and P-body-proteins Me31b and Patr-1 (Fig. 4G-N). Moreover, inhibiting the 26S proteasome by expression of dominant negative temperature sensitive proteasome β6 and β2 catalytic subunits (UAS-DTS-5,DTS-7), resulted in ectopic accumilaiton of Rogue and p-body proteins in ECs (Figs. 4O-R). Collectively suggesting that in young ECs p-body proteins are continuously ubiquitinated by the Kdo/CTLH complex and subsequently degraded by the 26S proteasome.

**Figure 4:**
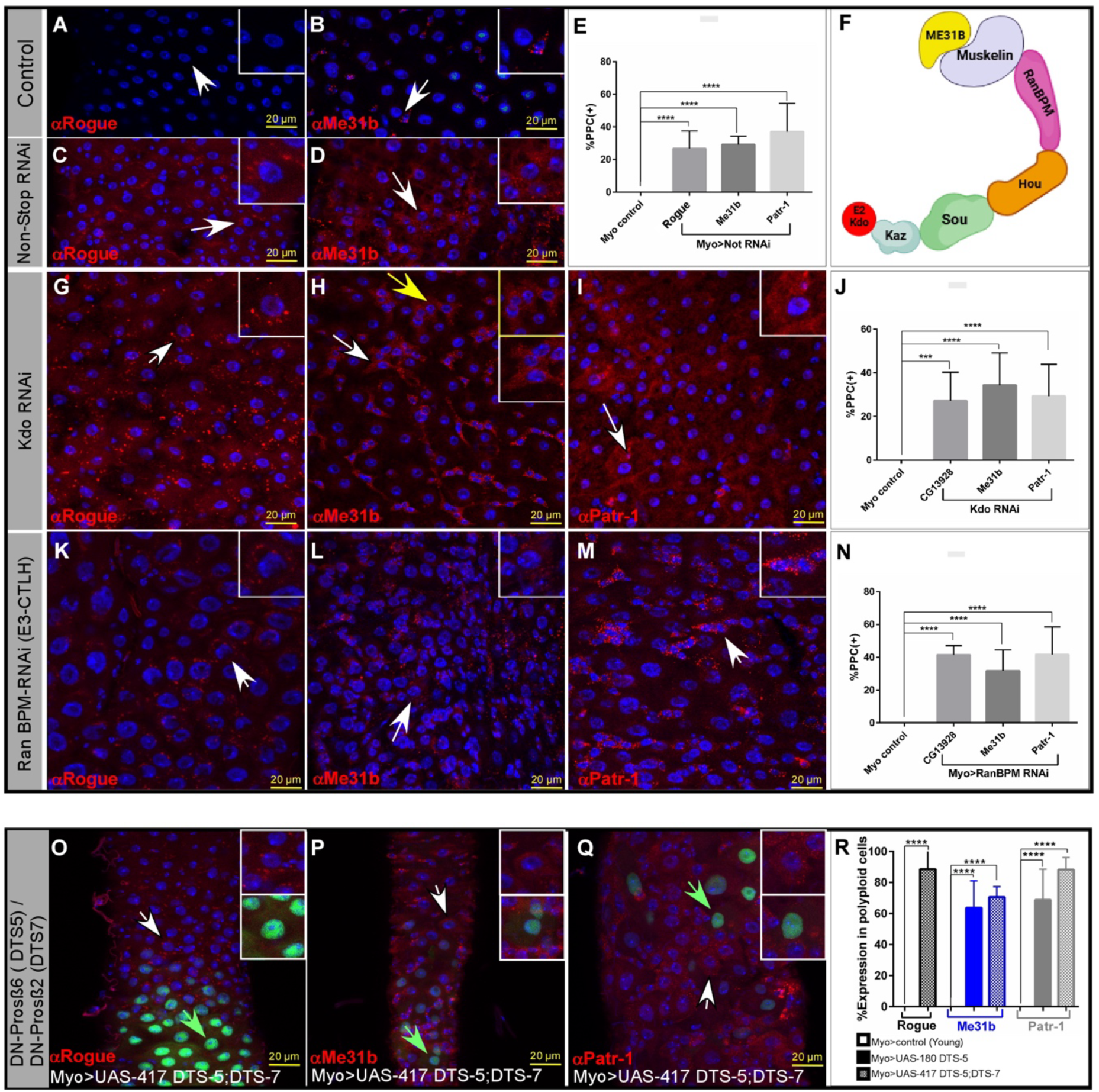
The DUB Non-stop along with the E2 Marie Kondo (Kdo) and the E3 Ub-ligase CTLH suppress the stem-cell∼related translational machinery. Confocal images and quantifications of mid-guts where ECs express the indicated transgenes, using the MyoIA>GAL4/ts80 system and immunostained with the indicated antibodies. DAPI marks DNA, Scale-bar is 20μM. **(A-E, G-J, K-N)** Elimination from young ECs of either Non-stop (C-E) or Kdo (G-J), or RanBPM (K-N), but not control (A, B), results in ectopic expression of Rogue (C, G, K), Me31B (D, H, L) or Patr-1 (I, M) proteins. **(F)** Diagram of Kdo∼CTLH (monomeric) ubiquitin ligase complex subunits and its substrate Me31b. **(O-R)** Expression of the indicated temperature sensitive dominant negative catalytic proteasome subunits β6, β2 (DN, DTS5, DTS7)^57, 58^ in ECs results in ectopic expression of p-body proteins ME31b and Patr1. The expression of these transgenes is partial, and therefore cells not expressing the transgenes maintain the Myo>GFP signal (green arrows), and do not express the indicated p-body proteins. Arrows point to cells in the insets. Quantification is shown in E, J, N, R and is based on three independent biological repeats ****=p<0.0001; ***=p<0.001.

Along these observations, elimination of Kdo using RNAi, but not control, from young ECs resulted in loss of EC identity as reflected in decline in EC proteins such as LamC, PDM1, and the junctional protein SSK (Fig. 5A, B, E-H, quantified in 5O; Fig.S7J, K). Concomitantly Kdo elimination resulted in ectopic expression of ISC related proteins such as Delta, LamDm0 (Figs. 5C, D, I, J and quantified in Fig. 5O). Moreover, expression of a FLAG-Kdo transgene in aged ECs suppressed the ectopic expression of Me31B (Fig. S7L-O). Likewise, eliminating the CTLH subunits RanBPM and Souji from young ECs resulted in loss of EC identity (e.g. reduced LamC protein levels and ectopic Delta expression; Figs. 5, K, L, M, N, and quantified in 5P). These observations suggest that in young ECs, the Kdo-CTLH complex safeguards ECbidentity by continuously eliminating the stem cell-related translational machinery including Rogue and P-body-proteins.

**Figure 5:**
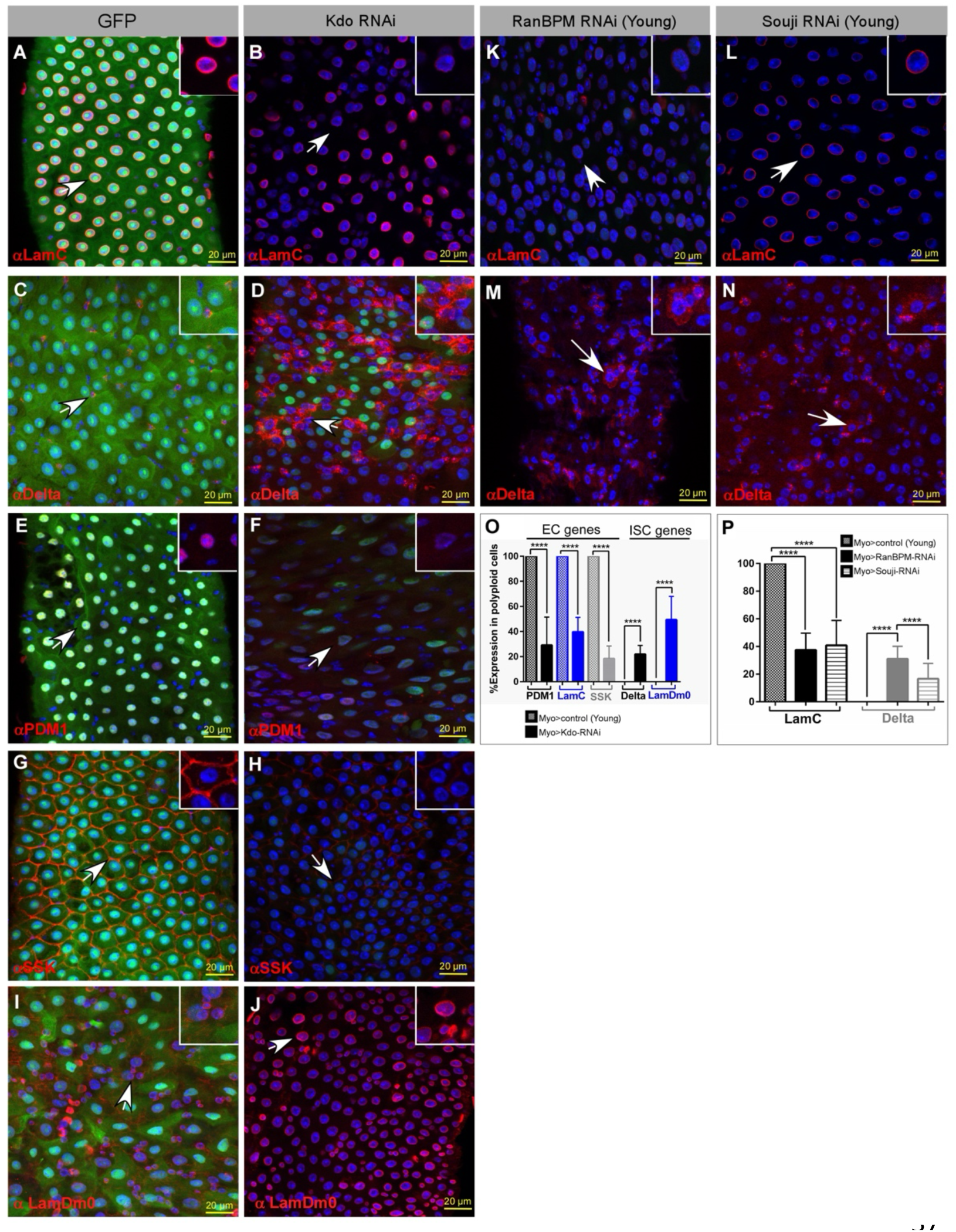
The Kdo∼CTLH E2/E3 complex supervise EC identity. Representative confocal images and quantifications of mid-guts where ECs express the indicated transgenes, using the MyoIA>GAL4/ts80 system and immunostained with the indicated antibodies. DAPI marks DNA, Scale-bar is 20μM. **(A-N)** Elimination from young ECs the E2 Kdo (B, D, F, H, J), or the E3 CTLH subunit RanBPM (K, M), or the CTLH subunit Souji (L, N), but not control (A, C, E, G, I.) results in reduced expression of LamC (B, K, L), Pdm1 (E, F), SSK (G, H), and concomitantly in ectopic expression Delta (C, D, M, N) and LmDm0 (I, J) along with reduction of the Myo::GFP signal. (O) and (P) are quantifications of three independent biological repeats and ****= p<0.0001.

### Rogue suppresses Kdo∼CTLH upon aging

Kdo and the CTLH complex subunits are present in young ECs (Figs. 6A-D, 6K). Remarkably, elimination of Kdo in young ECs results in reduced protein levels of multiple CTLH subunits including RanBPM, Musk, and Kaz (Fig. 6E-H and quantified in I). Suggesting that beyond its catalytic activity Kdo is a stabilizer of the CTLH complex. At the organismal level reducing Kdo levels in ECs shortened mean overall survival from 39 days to 32 days, a reduction of 18% in longevity (Fig. 6J). Previously it was shown that ME31b binds to Kdo mRNA and inhibits its translation^43^. We therefore hypothesized that the translational machinery driven by Rogue, and that pathologically accumulates upon aging, inactivates Kdo∼CTLH. Indeed, the proteins levels of Kdo decline upon aging, and elimination of Rogue in aged ECs restored Kdo protein levels (Fig. 6K-M and quantified in 6N).

**Figure 6:**
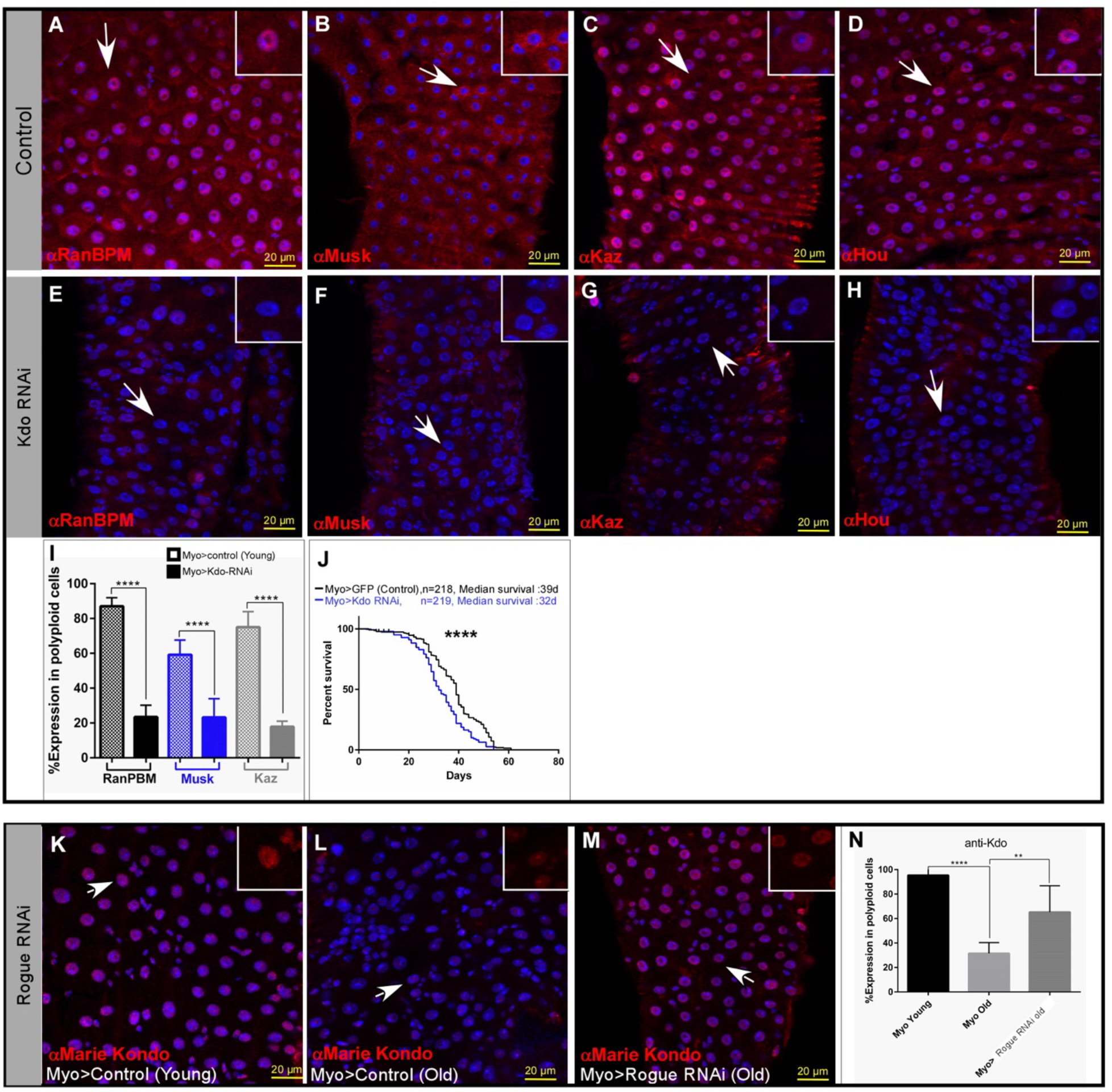
The E2 Kdo is required for the stability of CTLH subunits and is negatively regulated by Rogue. Confocal images of mid-guts where young ECs express control or UAS-Kdo RNAi transgenes genes, using the MyoIA-GAL4ts80 system and immunostained with the indicated antibodies against CTLH subunit proteins. White arrows point to cells shown in the inset, DAPI marks DNA, and Scale-bar is 20μM. **(A-D)** UAS-control, **(E-H)** UAS-Kdo RNAi. (I) Quantification of two independent experiments for each setting and ****p< 0.0001. **(J)** Elimination of Kdo using UAS-Kdo RNAi but not control from ECs reduces overall survival in ∼18%, ****=p<0.0001 and ***P<0.001. **(K-N)** Kdo protein is present in young EC (K), and its levels are reduced upon aging (L). Elimination of Rogue significantly restores Kdo protein levels in aged ECs (M) restored Kdo protein expression. Quantification is shown in (N), ****= P<0.0001, **=P<0.01, n=3.

Along these lines we observed that CTLH subunits are present in young ECs and forced expression of Rogue in young ECs resulted in reduced protein levels of CTLH complex subunits (Fig. 7A-H, quantified in O). Like in the case of Kdo, the levels of CTLH subunits declines upon aging, and elimination of Rogue from aged ECs using RNAi but not control, restored the protein level of CTLH subunits RanBPM, Hou, Muskelin, and to a lesser extent of Kaz (Figs. 7I-O). Most strikingly, manipulating Rogue levels in ECs affects overall animal survival: reducing Rogue levels via RNAi in ECs extended longevity from median of 39 to 48 days (∼23% increase in longevity), while its expression in young ECs reduced survival to 31 days (∼20% decrease in longevity; Fig. 5P). Thus, suggesting that the Kdo-CTLH complex supervises EC identity in young, differentiated EC, and that Kdo∼CTLH is cleared by Rogue upon aging.

**Figure 7:**
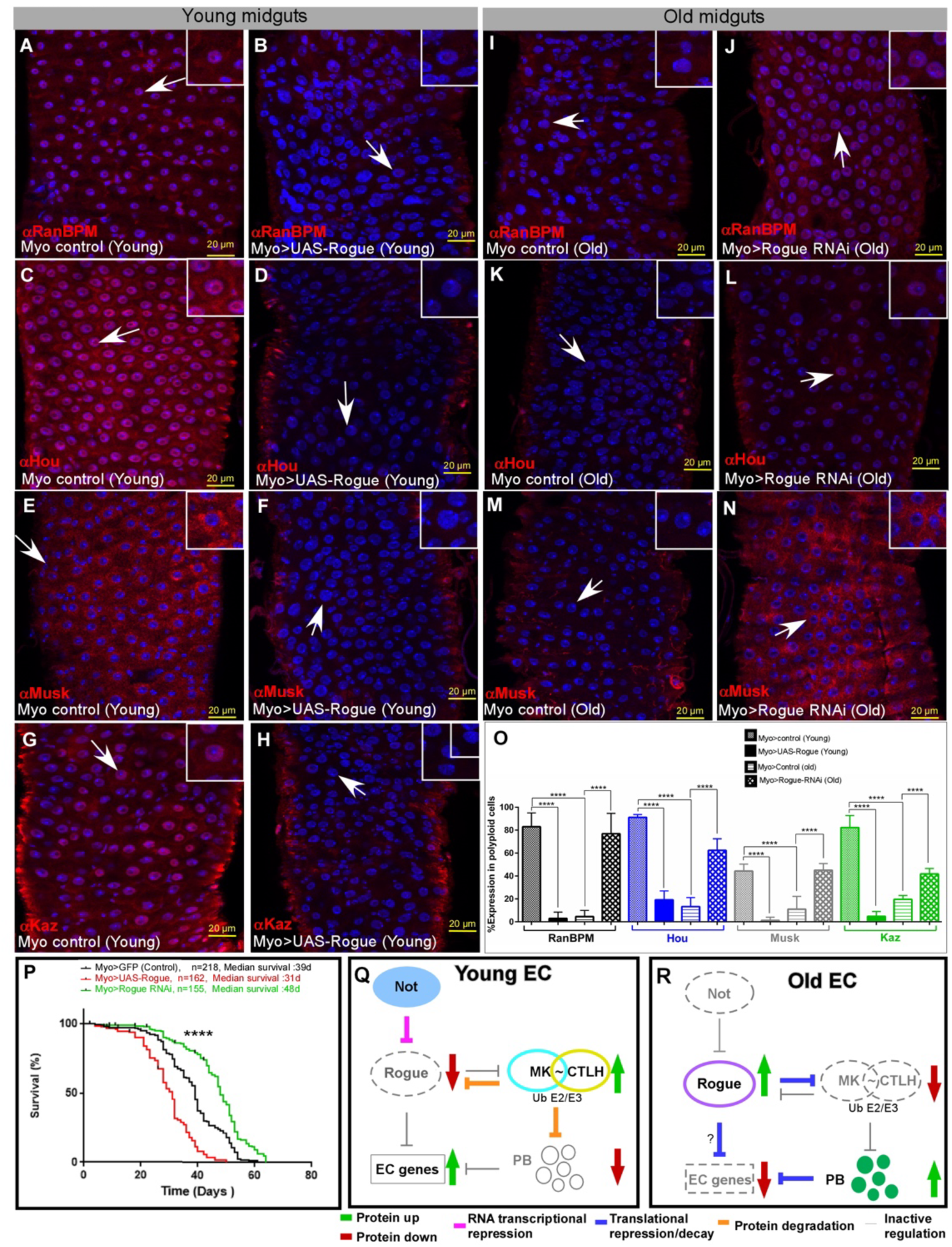
Rogue suppresses Kdo and CTLH proteins. **(A-N)** Representative confocal images of mid-guts where ECs express the indicated transgenes genes, using the MyoIA-GAL4ts80 system and immunostained with the indicated antibodies. DAPI marks DNA, Scale-bar is 20μM. (A-H) Expression of UAS-Rogue but not control in young ECs reduces the protein levels of RanBPM (A, B), Hou (C, D), Muskelin (E, F) and Kaz (G, H). (I-N) The level RanBPM, Hou and Muskelin declines in five weeks aged ECs (I, K, M), and is restored by the elimination of Rogue using UAS-Rogue RNAi (J, L, N). **(O)** Quantification of the above experiments of 3 independent biological repeats for each setting. **(P)** Elimination of Rogue from ECs using UAS-Rogue (CG13928)-RNAi extend overall longevity, and expression of UAS-Flag-Rogue in ECs reduces overall survival. Conditional expression in ECs was performed using the MyoIA-GAL4ts80 system. ****=p<0.0001. **(Q, R) A model for unlocking cell identity in aged EC by a Rogue-dependent translational machinery. (Q)** In young ECs, Non-stop suppresses *Rogue* mRNA. The E2 Kdo and CTLH E3 ligase complex reduce the protein levels of Rogue (if exist) and key P-body-proteins, safeguarding EC identity**. (R)** In aged ECs, the protein levels of Non-stop decline and Not-mediated repression of *Rogue* is canceled, resulting in ectopic expression of Rogue. Subsequently, the Kdo/CTLH E2/E3 are repressed by Rogue leading to re-appearance of P-body proteins and reactivation of stem-cell related translational repression machinery. The ectopic expression of P-bodies in aged ECs, inhibits EC gene signatures leading to loss of EC identity, see text for details.

## Discussion

### A model for post-transcriptional regulation of EC identity

Our experiments suggest a framework for post-transcriptional regulation of EC identity by antagonizing forces; the Kdo-CTLH E2/E3 ubiquitin ligase enzymes safeguard EC identity, and a stem cell-related translational machinery that is revived upon aging cancels EC identity (Figs. 7Q, R); In young EC, the DUB non-stop transcriptionally represses the mRNA expression of the translational repressor *Rogue*. In addition, the Kdo∼CTLH complex mediate the proteasomal elimination of P-body-proteins, and likely the minimal amounts of Rogue if present, altogether safeguarding EC identity. Upon aging, non-stop levels decline, and Rogue mRNA and protein levels accumulate (this study and ^16^). The ectopic expression of Rogue in ECs leads to the suppression of the ubiquitin E2/E3 Kdo-CTLH, re-appearance of the P-body machinery, and subsequent loss of EC identity.

### The delicate nature of the differentiated state

Upon aging, ECs that are no longer differentiated (PPC**) ectopically express multiple (>200) genes, suggesting that a key feature of identity maintenance is the silencing of previous and non-relevant fate genes. Some of these genes are potential IBs with diverse molecular functions and categories. Remarkably, in several cases, it is sufficient to suppress the expression of a single IB to restore EC identity, indicating that the balance that maintains EC identity is delicate. Yet, regardless of its origin (e.g. aging or genetic perturbations), loss of EC identity has a shared molecular framework that results in reversion to a “stem-cell like” organization. These changes include re-organization of the nuclear lamina (replacement of the differentiated LamC with ISC-related LamDm0), the ectopic expression of stem-cell related P-body-proteins, and reshaping the cell surface adhesion molecule replacing EC-adhesion proteins and ectopic with stem cell-related ligand adhesion molecules and (Delta and Arm, respectively). However, while PPC** (present either upon aging or expression of Rogue) ectopically express ISC genes, they do not revert to a functional stem-cells, nor do they trans-differentiate into EEs or other gut cell types. Suggesting that additional barriers for fate transition are still present. PPC** also maintain their polyploid state without entering mitosis or amitosis^46^. In turn and by co-expressing previous and non-relevant gene programs they enter a chaotic cell state that is detrimental, resulting in mis-differentiation of ISCs, loss of tissue homeostasis, impaired gut integrity, all resulting in accelerated aging and reduced animal survival. Furthermore, the observation that the elimination of a single IBs from ECs protects differentiated state, suppress aging phenotypes and extends longevity suggests that inhibition of IBs may have implication for therapy in aging-related disease.

In the *Drosophila* midgut we find that Kdo/CTLH regulation of the differentiated cell state is specific to ECs, but not to EEs the other differentiated cell type of the gut. Rogue or p-body proteins are not present in young or aged EEs, and elimination of Kdo from EE does not result in loss of EE identity. In this regard, it is worth noting that the nuclear organization of EEs is different from that of ECs; LamDm0 is the dominant EE nuclear Lamin, similar to ISCs, and not LamC as in the case of ECs. Suggesting that EEs maintain their differentiated state using a yet to be discovered mechanism(s).

### Kdo∼CTLH maintain differentiated cell identity

Conserved from yeast (termed GID) to humans, the Kdo∼CTLH RING-type macromolecular E2/E3 complex regulates a plethora of biological processes including embryogenesis, erythroid differentiation, mTor signaling, metabolism, immune response, and cancer^47–55^. In *Drosophila,* during embryogenesis, Kdo/CTLH complex is required to ubiquitylate maternally deposited P-body-proteins such as Me31B, TRAL and Patr-1, leading to their proteasomal degradation^43,44^. While differentiation was extensively studied at the gene-expression level for decades, it is now clear that remodeling the cell’s proteome is not less important. In this regard, it was recently shown that CTLH-dependent ubiquitylation and degradation of earlier fate proteins govern erythroid differentiation^47,55^. Here we discovered that in the adult midgut, the Kdo∼CTLH complex is critical not only to differentiation per-se but is critical to maintain the differentiated state by the continuous degradation of the stem cell-related translational machinery in young, differentiated ECs. Suggesting that the role of CTLH in targeting the translational machinery/RNA binding proteins is a universal principle from flies to humans. Interestingly, the human ortholog of Non-stop, USP22, was recently shown to be a barrier for iPS reprograming^14^ and is pro-tumorigenic in RAS-driven leukemia^59^ further demonstrating the role of conserved IS as molecular machinery preventing fate transition and tumorigenesis

It is not fully clear how the CTLH complex recognizes its diverse substrates, likely involving more than one substrate recognition subunit^53^. Recently, the CTLH subunit Muskelin was shown to be the substrate recognition subunit for these RNA-binding proteins during embryogenesis such as Me31B^56^. Indeed, in the midgut, loss of Muskelin resulted in increased levels of Me31B and TRAL. In contrast, the fly ortholog of WDR26, CG7611, which is also suggested to be a recognition subunit of the CTLH complex^53^ and is mutually exclusive with Muskelin^56^, is not involved in maintaining EC identity (not shown). The regulation of Kdo∼CTLH complex itself is predominantly post-transcriptional. Unlike the case of erythropoiesis where the mRNA levels of the E2 UbCH8 increase upon erythroblast differentiation^47,55^, the mRNA levels of the E2 Kdo as well as subunits of the CTLH complex do not change upon differentiation of ISC to EC, nor do the mRNAs of these proteins decline upon aging. One exception is the subunit Yippee (CG1989, ortholog of YPEL5) whose mRNA expression increased upon progenitor differentiation into ECs (Fig.S9L), yet the importance of this observation is currently not clear.

### Reactivation of a stem cell-related translation repression machinery upon aging

The levels of the identity supervisors Non-stop and the CTLH complex subunit proteins sharply decline in aged ECs (this study and ^16^). The decline in CTLH is attributed to ectopic expression of Rogue and reactivation of the P-body machinery. The exact mechanism(s) are not fully understood, and may be due, at least in part, to the rise of Me31B protein that was shown to bind to Kdo mRNA directly and inhibits its translation^43^. This rise is observed physiologically in aged ECs, expression of Rogue, or Rogue accumulation upon loss of the identity supervisor non-stop in young EC. The loss of Kdo is pivotal beyond catalyzing the ubiquitination of CTLH substrates, as its loss results in clearance of multiple CTLH subunits. In addition, the association of Rogue with 40-60S monosome fractions, likely has significant implication on overall translation re-shaping the EC proteome^42^.

To conclude, herein we discovered that maintaining the differentiated state of enterocytes requires the continuous degradation of a stem-cell related translational repression machinery by a dedicated ubiquitin E2/E3 pair. This regulation that declines upon aging leading, unlocking the differentiated state, impairs tissue homeostasis and limits overall survival.

## Acknowledgments

We are grateful to Nickolas Sokol, Kausik Si, Bruce Edgar, Hermann Steller and Norbert Perrimon for sharing antibodies and fly lines. We thank Alan Schwartz, Dan Finley, Miguel Prado, and Brenda Schulman for discussions and insightful comments.

## Author contributions

Conceptualization: SD, EBL, AO, RDM

Investigation: SD, RG, EBL, MM JS, JH

Visualization: SD, EBL, MM

Funding acquisition: OSR, RMD, AS, AO

Supervision: OSR, RMD, AS, AO

Writing – original draft: SD, ELB, MM, AO

Writing – review & editing: SD, ELB, MM, RMD, OSR, AO

## Data and materials availability

Sc-RNA seq data is available at: GEO submission # GSE289519

All other data are available in the main text or the supplementary materials.

## Supplementary Figures Materials and methods

Supplemental figures S1-S8 and legend,

Table S1 expression signatures,

Materials and Methods include:

- Key resource table with fly stocks and antibodies used in this study
- Plasmids and Primers used in this study
- Chemicals used

*Methods*

- Generation of UAS-HA-FLAG-Kdo transgenic line.
- Conditional expression of transgenes in specific gut cells
- Conditional G-TRACE analysis
- Gut dissection and immunofluorescence detection
- Gut integrity and tracing of organismal survival
- Genomic analysis; GTACE-coupled sc-RNAseq and bioinformatics analyses.
- Polysome profiling
- Statistical analyses

## References

1. Blau, H.M., and Baltimore, D. (1991). Differentiation requires continuous regulation. The Journal of Cell Biology 112, 781. 10.1083/JCB.112.5.781.

2. Sánchez Alvarado, A., and Yamanaka, S. (2014). Rethinking differentiation: Stem cells, regeneration, and plasticity. Cell 157, 110–119. 10.1016/J.CELL.2014.02.041/ASSET/237F77F1-26C3-410D-9E28-F043F4FC56E5/MAIN.ASSETS/GR5.JPG.

3. Natoli, G. (2010). Maintaining cell identity through global control of genomic organization. Immunity 33, 12–24. 10.1016/J.IMMUNI.2010.07.006/ASSET/6E145666-AE00-43FC-81C1-F58375B331E1/MAIN.ASSETS/GR4.JPG.

4. Holmberg, J., and Perlmann, T. (2012). Maintaining differentiated cellular identity. Nature Reviews Genetics 2012 13:6 13, 429–439. 10.1038/nrg3209.

5. Fazilaty, H., and Basler, K. (2023). Reactivation of embryonic genetic programs in tissue regeneration and disease. Nat Genet 55, 1792–1806. 10.1038/s41588-023-01526-4.

6. Bitman-Lotan, E., and Orian, A. (2021). Nuclear organization and regulation of the differentiated state. Cell. Mol. Life Sci. 78, 3141–3158. 10.1007/s00018-020-03731-4.

7. Deneris, E.S., and Hobert, O. (2014). Maintenance of postmitotic neuronal cell identity. Nat Neurosci 17, 899–907. 10.1038/nn.3731.

8. Ocampo, A., Reddy, P., and Belmonte, J.C.I. (2016). Anti-Aging Strategies Based on Cellular Reprogramming. Trends in Molecular Medicine 22, 725–738. 10.1016/j.molmed.2016.06.005.

9. Mikkola, I., Heavey, B., Horcher, M., and Busslinger, M. (2002). Reversion of B cell commitment upon loss of Pax5 expression. Science 297, 110–113. 10.1126/SCIENCE.1067518.

10. Cobaleda, C., Jochum, W., and Busslinger, M. (2007). Conversion of mature B cells into T cells by dedifferentiation to uncommitted progenitors. Nature 449, 473–477. 10.1038/nature06159.

11. Schwitalla, S., Fingerle, A.A., Cammareri, P., Nebelsiek, T., Göktuna, S.I., Ziegler, P.K., Canli, O., Heijmans, J., Huels, D.J., Moreaux, G., et al. (2013). Intestinal Tumorigenesis Initiated by Dedifferentiation and Acquisition of Stem-Cell-like Properties. Cell 152, 25–38. 10.1016/j.cell.2012.12.012.

12. Hanahan, D. (2022). Hallmarks of Cancer: New Dimensions. Cancer Discovery 12, 31–46. 10.1158/2159-8290.CD-21-1059.

13. Qin, H., Diaz, A., Blouin, L., Lebbink, R.J., Patena, W., Tanbun, P., LeProust, E.M., McManus, M.T., Song, J.S., and Ramalho-Santos, M. (2014). Systematic Identification of Barriers to Human iPSC Generation. Cell 158, 449–461. 10.1016/j.cell.2014.05.040.

14. Gürhan, G., Sevinç, K., Aztekin, C., Gayretli, M., Yılmaz, A., Yıldız, A.B., Ervatan, E.N., Morova, T., Datlı, E., Coleman, O.D., et al. (2025). A chromatin-focused CRISPR screen identifies USP22 as a barrier to somatic cell reprogramming. Communications biology 8, 454. 10.1038/S42003-025-07899-Y.

15. McCarthy, R.L., Zhang, J., and Zaret, K.S. (2023). Diverse heterochromatin states restricting cell identity and reprogramming. Trends in Biochemical Sciences 48, 513–526. 10.1016/j.tibs.2023.02.007.

16. Erez, N., Israitel, L., Bitman-Lotan, E., Wong, W.H., Raz, G., Cornelio-Parra, D.V., Danial, S., Flint Brodsly, N., Belova, E., Maksimenko, O., et al. (2021). A Non-stop identity complex (NIC) supervises enterocyte identity and protects from premature aging. eLife 10, e62312. 10.7554/eLife.62312.

17. Flint Brodsly, N., Bitman-Lotan, E., Boico, O., Shafat, A., Monastirioti, M., Gessler, M., Delidakis, C., Rincon-Arano, H., and Orian, A. (2019). The transcription factor Hey and nuclear lamins specify and maintain cell identity. eLife 8, e44745. 10.7554/eLife.44745.

18. Weintraub, H., Tapscott, S.J., Davis, R.L., Thayer, M.J., Adam, M.A., Lassar, A.B., and Miller, A.D. (1989). Activation of muscle-specific genes in pigment, nerve, fat, liver, and fibroblast cell lines by forced expression of MyoD. Proceedings of the National Academy of Sciences of the United States of America 86, 5434–5438. 10.1073/PNAS.86.14.5434.

19. Dall’Agnese, A., Caputo, L., Nicoletti, C., Di Iulio, J., Schmitt, A., Gatto, S., Diao, Y., Ye, Z., Forcato, M., Perera, R., et al. (2019). Transcription Factor-Directed Re-wiring of Chromatin Architecture for Somatic Cell Nuclear Reprogramming toward trans-Differentiation. Molecular Cell 76, 453–472.e8. 10.1016/j.molcel.2019.07.036.

20. Takahashi, K., and Yamanaka, S. (2006). Induction of pluripotent stem cells from mouse embryonic and adult fibroblast cultures by defined factors. Cell 126, 663–676. 10.1016/j.cell.2006.07.024.

21. Müller, M., Hermann, P.C., Liebau, S., Weidgang, C., Seufferlein, T., Kleger, A., and Perkhofer, L. (2016). The role of pluripotency factors to drive stemness in gastrointestinal cancer. Stem Cell Research 16, 349–357. 10.1016/j.scr.2016.02.005.

22. Jasper H. (2020) Intestinal Stem Cell Aging: Origins and Interventions. Annu Rev Physiol. 2020 Feb 10;82:203-226. doi: 10.1146/annurev-physiol-021119-034359.

23. Lemaitre, B., and Miguel-Aliaga, I. (2013). The Digestive Tract of *Drosophila melanogaster*. Annu. Rev. Genet. 47, 377–404. 10.1146/annurev-genet-111212-133343.

24. Jiang, H., and Edgar, B.A. (2012). Intestinal stem cell function in Drosophila and mice. Current Opinion in Genetics & Development 22, 354–360. 10.1016/j.gde.2012.04.002.

25. Zhang, P., and Edgar, B.A. (2022). Insect Gut Regeneration. Cold Spring Harb Perspect Biol 14, a040915. 10.1101/cshperspect.a040915.

26. Micchelli, C.A., and Perrimon, N. (2006). Evidence that stem cells reside in the adult Drosophila midgut epithelium. Nature 439, 475–479. 10.1038/nature04371.

27. Buchon, N., Osman, D., David, F.P.A., Yu Fang, H., Boquete, J.-P., Deplancke, B., and Lemaitre, B. (2013). Morphological and Molecular Characterization of Adult Midgut Compartmentalization in Drosophila. Cell Reports 3, 1725–1738. 10.1016/j.celrep.2013.04.001.

28. Marianes, A., and Spradling, A.C. (2013). Physiological and stem cell compartmentalization within the Drosophila midgut. eLife 2, e00886. 10.7554/eLife.00886.

29. Becker, J.S., Mccarthy, R.L., Sidoli, S., Lin, S., Garcia, B.A., and Zaret Correspondence, K.S. (2017). Genomic and proteomic resolution of heterochromatin and its restriction of alternate fate genes. cell.comJS Becker, RL McCarthy, S Sidoli, G Donahue, KE Kaeding, Z He, S Lin, BA Garcia, KS ZaretMolecular cell, 2017•cell.com1023-1037, 68 .e15. 10.1016/j.molcel.2017.11.030.

30. Chen, H., Zheng, X., and Zheng, Y. (2015). Lamin-B in systemic inflammation, tissue homeostasis, and aging. Nucleus 6, 183–186. 10.1080/19491034.2015.1040212.

31. Evans, C.J., Olson, J.M., Ngo, K.T., Kim, E., Lee, N.E., Kuoy, E., Patananan, A.N., Sitz, D., Tran, P., Do, M.-T., et al. (2009). G-TRACE: rapid Gal4-based cell lineage analysis in Drosophila. Nat Methods 6, 603–605. 10.1038/nmeth.1356.

32. Hung, R.-J., Hu, Y., Kirchner, R., Liu, Y., Xu, C., Comjean, A., Tattikota, S.G., Li, F., Song, W., Ho Sui, S., et al. (2020). A cell atlas of the adult Drosophila midgut. Proc. Natl. Acad. Sci. U.S.A. 117, 1514–1523. 10.1073/pnas.1916820117.

33. Li, H., Qi, Y., and Jasper, H. (2016). Preventing Age-Related Decline of Gut Compartmentalization Limits Microbiota Dysbiosis and Extends Lifespan. Cell Host & Microbe 19, 240–253. 10.1016/j.chom.2016.01.008.

34. Jasper, H. (2020). Intestinal Stem Cell Aging: Origins and Interventions. Annual Review of Physiology 82, 203–226. 10.1146/ANNUREV-PHYSIOL-021119-034359.

35. Khan, M.R., Li, L., Pérez-Sánchez, C., Saraf, A., Florens, L., Slaughter, B.D., Unruh, J.R., and Si, K. (2015). Amyloidogenic Oligomerization Transforms Drosophila Orb2 from a Translation Repressor to an Activator. Cell 163, 1468–1483. 10.1016/j.cell.2015.11.020.

36. Chen, X., and Dickman, D. (2021). Tissue-Specific Ribosome Profiling in Drosophila. Methods in Molecular Biology 2252, 175–188. 10.1007/978-1-0716-1150-0_7.

37. Bar-Yosef, H., Venezian, J., Klann, K., and Shiber, A. (2022). Purification of Ribosome-Nascent-Chain Complex for Ribosome Profiling and Selective Ribosome Profiling. Methods in Molecular Biology 2477, 179–193. 10.1007/978-1-0716-2257-5_11.

38. Jazdowska-Zagrodzińska, B. (1966). Experimental studies on the role of “polar granules” in the segregation of pole cells in Drosophila melanogaster. J Embryol Exp Morphol 16, 391–399.

39. Updike, D.L., Kek Upa’a Knutson, A., Egelhofer, T.A., Campbell, A.C., and Strome, S. (2014). Germ-granule components prevent somatic development in the C. elegans germline. cell.comDL Updike, AK Knutson, TA Egelhofer, AC Campbell, S StromeCurrent Biology, 2014•cell.com975–970, 24. 10.1016/j.cub.2014.03.015.

40. Trcek, T., and Lehmann, R. (2019). Germ granules in Drosophila. Traffic 20, 650–660. 10.1111/TRA.12674.

41. Buddika, K., Huang, Y.-T., Ariyapala, I.S., Butrum-Griffith, A., Norrell, S.A., O’Connor, A.M., Patel, V.K., Rector, S.A., Slovan, M., Sokolowski, M., et al. (2022). Coordinated repression of pro-differentiation genes via P-bodies and transcription maintains Drosophila intestinal stem cell identity. Current Biology 32, 386–397.e6. 10.1016/j.cub.2021.11.032.

42. Smith, P.R., Loerch, S., Kunder, N., Stanowick, A.D., Lou, T.-F., and Campbell, Z.T. Functionally distinct roles for eEF2K in the control of ribosome availability and p-body abundance. nature.comPR Smith, S Loerch, N Kunder, AD Stanowick, TF Lou, ZT CampbellNature communications, 2021•nature.com. 10.1038/s41467-021-27160-4.

43. Zavortink, M., Rutt, L.N., Dzitoyeva, S., Henriksen, J.C., Barrington, C., Bilodeau, D.Y., Wang, M., Chen, X.X.L., and Rissland, O.S. (2020). The E2 Marie Kondo and the CTLH E3 ligase clear deposited RNA binding proteins during the maternal-to-zygotic transition. eLife 9, e53889. 10.7554/eLife.53889.

44. Cao, W.X., Kabelitz, S., Gupta, M., Yeung, E., Lin, S., Rammelt, C., Ihling, C., Pekovic, F., Low, T.C.H., Siddiqui, N.U., et al. (2020). Precise Temporal Regulation of Post-transcriptional Repressors Is Required for an Orderly Drosophila Maternal-to-Zygotic Transition. Cell Reports 31, 107783. 10.1016/j.celrep.2020.107783.

45. Liu, H., and Pfirrmann, T. (2019). The Gid-complex: an emerging player in the ubiquitin ligase league. Biological Chemistry 400, 1429–1441. 10.1515/hsz-2019-0139.

46. Lucchetta, E.M., and Ohlstein, B. (2017). Amitosis of Polyploid Cells Regenerates Functional Stem Cells in the Drosophila Intestine. Cell Stem Cell 20, 609–620.e6. 10.1016/j.stem.2017.02.012.

47. Sherpa, D., Mueller, J., Karayel, Ö., Xu, P., Yao, Y., Chrustowicz, J., Gottemukkala, K.V., Baumann, C., Gross, A., Czarnecki, O., et al. (2022). Modular UBE2H-CTLH E2-E3 complexes regulate erythroid maturation. eLife 11, e77937. 10.7554/eLife.77937.

48. Santt, O., Pfirrmann, T., Braun, B., Juretschke, J., Kimmig, P., Scheel, H., Hofmann, K., Thumm, M., and Wolf, D.H. (2008). The yeast GID complex, a novel ubiquitin ligase (E3) involved in the regulation of carbohydrate metabolism. Molecular Biology of the Cell 19, 3323–3333. 10.1091/MBC.E08-03-0328.

49. Simwela, N.V., Johnston, L., Bitar, P.P., Jaecklein, E., Altier, C., Sassetti, C.M., and Russell, D.G. (2024). Genome-wide screen of Mycobacterium tuberculosis-infected macrophages revealed GID/CTLH complex-mediated modulation of bacterial growth. Nat Commun 15, 9322. 10.1038/s41467-024-53637-z.

50. Mctavish, C.J., Bérubé-Janzen, W., Wang, X., Maitland, M.E.R., Salemi, L.M., Hess, D.A., and Schild-Poulter, C. Regulation of c-Raf Stability through the CTLH Complex. mdpi.comCJ McTavish, W Bérubé-Janzen, X Wang, MER Maitland, LM Salemi, DA HessInternational journal of molecular sciences, 2019•mdpi.com. 10.3390/ijms20040934.

51. Salemi, L.M., Maitland, M.E.R., McTavish, C.J., and Schild-Poulter, C. (2017). Cell signalling pathway regulation by RanBPM: molecular insights and disease implications. Open Biol. 7, 170081. 10.1098/rsob.170081.

52. Qiao, S., Langlois, C.R., Chrustowicz, J., Sherpa, D., Karayel, O., Hansen, F.M., Beier, V., Von Gronau, S., Bollschweiler, D., Schäfer, T., et al. (2020). Interconversion between Anticipatory and Active GID E3 Ubiquitin Ligase Conformations via Metabolically Driven Substrate Receptor Assembly. Molecular Cell 77, 150–163.e9. 10.1016/j.molcel.2019.10.009.

53. Gottemukkala, K.V., Chrustowicz, J., Sherpa, D., Sepic, S., Vu, D.T., Karayel, Ö., Papadopoulou, E.C., Gross, A., Schorpp, K., von Gronau, S., et al. (2024). Non-canonical substrate recognition by the human WDR26-CTLH E3 ligase regulates prodrug metabolism. Molecular Cell 84, 1948–1963.e11. 10.1016/J.MOLCEL.2024.04.014/ASSET/F45DE557-CB39-40A9-BB61-45342EEAF86E/MAIN.ASSETS/FX1_LRG.JPG.

54. Yi, S.A., Sepic, S., Schulman, B.A., Ordureau, A., and An, H. (2024). mTORC1-CTLH E3 ligase regulates the degradation of HMG-CoA synthase 1 through the Pro/N-degron pathway. Molecular Cell 84, 2166–2184.e9. 10.1016/j.molcel.2024.04.026.

55. Ren, K., Li, E., and Ji, P. (2022). Proteome remodeling and organelle clearance in mammalian terminal erythropoiesis. Current Opinion in Hematology 29, 137–143. 10.1097/MOH.0000000000000707

56. Briney, C.A., Henriksen, J.C., Lin, C., Jones, L.A., Benner, L., Rains, A.B., Gutierrez, R., Gafken, P.R., and Rissland, O.S. (2025). Muskelin is a substrate adaptor of the highly regulated Drosophila embryonic CTLH E3 ligase. EMBO Rep 26, 1647–1669. 10.1038/s44319-025-00397-6.

57. Belote JM, Fortier E. Targeted expression of dominant negative proteasome mutants in Drosophila melanogaster. Genesis. 2002 34 :80–2. doi: 10.1002/gene.10131.

58. Bader M, Benjamin S, Wapinski OL, Smith DM, Goldberg AL, Steller H. (2011) A conserved F box regulatory complex controls proteasome activity in *Drosophila*. Cell. 145 :371–82

59. Melo-Cardenas J, Xu Y, Wei J, Tan C, Kong S, Gao B, Montauti E, Kirsammer G, Licht JD, Yu J, Ji P, Crispino JD, Fang D. (2018) USP22 deficiency leads to myeloid leukemia upon oncogenic Kras activation through a PU.1-dependent mechanism. Blood. 2132:423-434. doi: 10.1182/blood-2017-10-811760.

