## supplemental figures and methods for "A Ubiquitin network safeguards cell identity by continuously degrading a stem-cell related translational machinery"

**Supplemental materials The S1 file includes:**

1. Materials and Methods
2. Legend to S1-S8
3. Tables S1 (Excell file)
4. Resource Table
5. References

### 1. Materials and Methods

**Supplementary Materials**
**Materials and Methods:**

- Key resource table with fly stocks and antibodies used in this study
- Plasmids and Primers used in this study
- Chemicals used

*Methods:*

- Generation of UAS-HA-FLAG-Kdo transgenic line.
- Conditional expression of transgenes in specific gut cells
- Conditional G-TRACE analysis
- Gut dissection and immunofluorescence detection
- Gut integrity and tracing of organismal survival
- Genomic analysis; Sc-RNA-seq and bioinformatics analyses.
- Polysome profiling
- Statistical analyses

**Fly stocks used in this study:** Fly stocks were maintained on yeast-cornmeal-molasses-malt extract medium at 18°C or as stated in the text.

**UAS and Gal4 transgenic lines used**: All lines used in this study are described in the Key Resources table.

**Antibodies used in this study**: All primary and secondary antibodies used are described in the Key resource table.

**Chemicals:** Bromophenol Blue (Sigma #B5525), Guanidine hydrochloride (Sigma #G4505), NP40 (Ipegal CA-630) (Sigma #I3021), Triton X-100 (Amresco #0694), Acrylamide (Bis-Acrylamide 29:1) (Biological Industries #01-874-1A), Ammonium Persulfate (Sigma #A-9164), TEMED (Sigma #T-7024), L-Glutamine (Gibco #25030024), MG132 (Boston Biochemicals), Blot Qualified BSA (Biological Industries #PRW3841), KCl, MgCl2, Heparin (Sigma #H4784), Bradford Protein Assay (BioRad #500-0006), EZ-ECL (Biological Industries #20-500-500), FD&C blue dye #1, Cyclohexamide (Sigma #01810), Elastase (Sigma-Aldrich, E0258), Collagenase type 3 (LS004182 - Worthington Biochemical cooperation), Collagenase type 4 (LS004188 – Worthington Biochemical cooperation).

***Methods:***

***Generation of g UAS-HA-FLAG-Kdo transgenic flies:***

A plasmid coding for CG2257 (Kdo) CDs was obtained from FlyBase and synthesized with either a N- or C-terminal HA-Flag2 tags into pUASt vector (Vector Builder Inc, Chicago, Il) . The plasmids were injected and integrated at chromosome 3 using thePhiC3 integrase system and parent line y[1] M{RFP[3xP3.PB] GFP[E.3xP3]=vas-int.Dm}ZH-2A w[*]; PBac{y[+]-attP-9A}VK00005[1] (Rainbow Transgenic Flies Inc. Camarillo, CA). Stable transformants were selected based on *miniwhite* expression and locus integration was confirmed by PCR.

***Conditional expression of transgenes in specific gut cells:*** Conditional expression of transgenic lines in specific midgut cells was achieved by activating a UAS-transgene under the expression of the indicated cell-specific Gal4-driver together with tub::Gal80$^{ts}$[2–4]. Flies were reared at 18°C. 4 days old, F1 adult females' progeny were transferred to the restrictive temperature 29°C (Gal80 off, Gal4 on) for one week unless indicated otherwise, dissected and analyzed. At least three biological independent repeats were performed for each experiment. Where possible, multiple RNAi lines were used. Similar results were observed in males.

***G-TRACE analysis*:** G-TRACE analyses was as described in[5–7] using either Myo-Gal4; G-TRACE young, or five weeks old flies. Were indicated these transgenes also expressed either UAS-GFP; Gal80$^{ts}$ (control) or UAS-Rogue-CG13928 RNAi; Gal80$^{ts}$ generating the appropriate genotypes. Files were raised at 18°C (a temperature where no G-TRACE signal was detected). At two to four days adult females were transferred to 29°C and linage tracing was performed.

***Gut dissection and immunofluorescence detection:*** Gut fixation and staining were carried out as previously described[4–6,8].

**G-TRACE coupled scRNA-seq profiling:** scRNA-seq was performed similar to[9]. *Drosophila* guts were dissociated to single cells. Guts were dissected from the following groups: 7-day old control or Rogue expressing UAS-transgenes, and 5 weeks old Myo-Gal4; G-TRACE adult transgenic female flies. From each group, 45 flies were dissected into ice cold PBS containing 1% BSA in dissection plates. After gut dissection, the crop and midgut/hindgut junction and Malpighian tubules were removed. Dissected guts were chopped into small pieces using a razorblade and immediately transferred to an Eppendorf tube containing 400µl of digestion solution: 4 mg/ml elastase solution (E0258), 50 mg/ml collagenase type 3 (LS004182) and 50 mg/ml collagenase type 4 (LS004188). Samples were vortexed for 20" and incubated on a shaker at 27°C for 30 minutes. The cell suspension was filtered through a 70µm cell strainer, and spined down at 100 x g for 5 min at 4°C. The cells pellet was resuspended in 400μl of cold PBS containing 1% BSA. 10μl of the resuspended cells was used to determine cell viability using 0.4% trypan blue staining and cells were counted using a hemocytometer. Subsequently the samples were subjected to single cell sequencing using 10x Genomics. Library preparation and data generation Two RNA single cell library was prepared according to 10X manufacture protocol (Chromium Next GEM Single Cell 3' Library & Gel Bead Kit v3.1, PN-1000268) using 12500 input cells for sample 'old' and 20000 input cells for sample 'young'. Single cell separation was performed using the Chromium Next GEM Chip G Single Cell Kit (PN-1000120). After construction, the concentration of library was measured using Qubit (Invitrogen) and the size was determined using the TapeStation 4200 with the High Sensitivity D1000 kit (cat no. 5067-5584). The RNAseq data was generated on Illumina NextSeq2000, P2 100 cycles (R1-28bp, R2-90bp, I1-10bp, I2-10bp) (Illumina, cat no. 20046811)

***Bioinformatic analysis**:* CellRanger pipeline (v6.0.1, 10x genomics) mkref command was used to create the *Drosophila Melanogaster* costume genome, BDGP6.28.102, with GFP and RFP sequences. Cellranger count with default parameters was used for alignment filtering, barcode counting, and UMI counting. This resulted in 2,279 and 1,789 cells for the old and young samples, respectively. The Seurat R package (v4.0.4) was used for downstream analysis and visualization. Gene-cell matrices were filtered to remove cells with less than 500 UMIs, less than 250 genes, and more than 25% of mitochondrial reads. After implementing these quality control measures, 1,436 young cells, xx young cells overexpressing Rogue and 1,789 old cells, and 12,434 genes were retained for further analysis.

The expression data was normalized using Seurat's NormalizeData function, which normalizes the feature expression measurements for each cell by the total expression, multiplies this by a scale factor (10,000), and then log-transforms the results. The young gut sample was first analyzed separately. The top 2,000 highly variable genes were identified using Seurat's "FindVariableFeatures" function with the 'vst' method. Potential sources of unspecific variation in the data were removed by regressing out the mitochondrial gene proportion and UMI count effect using linear models and finally by scaling and centering the residuals as implemented in the function "ScaleData" of the Seurat package. Principal component analysis (PCA) was performed. We selected 25 principal components (PC) for downstream analyses. Cell clusters were generated using Seurat's unsupervised graph-based clustering functions "FindNeighbors" and "FindClusters" (resolution = 0.5). UMAP was generated using the "RunUMAP" on the projected principal component (PC) space. Clusters were annotated manually using specific markers expression. The old midgut sample was then integrated with the young sample using Seurat's integration functions[3]. Briefly, the "FindIntegrationAnchors" function was used to identify anchors between the two samples, and the "MapQuery" function was used to annotate the old sample cells and add UMAP reduction coordinates for each cell. Cells were assigned as RFP or GFP positive if the expression of these genes were at least 1. DE analysis between different groups of cells was done using the "FindMarkers" function with default parameters (non-parametric Wilcoxon rank sum test). Seurat's functions FeaturePlot and DimPlot were used for visualization. Seurat's DotPlot and VlnPlot function were used to visualize gene expression for specific groups. Plots were further formatted using custom R scripts with the packages ggplot2 [10–14].

**Polysome profiling:** For polysome profiling 150 adult female flies of the following genotypes were used: 1. 7-day old Mex-Gal4>UAS-GFP control, 2. 7-day old Mex-Gal4>UAS- Rogue (CG13928)-FLAG. 3. Five weeks old expressing Mex-Gal4>UAS-GFP (control), or 4. Five weeks old expressing Mex-Gal4>UAS- Rogue CG13928-RNAi. The Mex-Gal4 drives the expression of UAS-transgenes solely in adult EC and no other cell in the entire adult fly (fly atlas. Polysome profiling was performed in three biological replicates. Flies were frozen in liquid nitrogen and heads, wings and limbs were removed, and abdomens were cryo-lysed in lysis buffer (20 mM Tris-HCl pH 8, 140 mM KCl, 10 mM MgCl2, 0.1 % NP-40 0.5mg/ml CHX, 1mg/ml Heparin) by tissue LyserII (Qiagen). Lysates were centrifuged at 13,000 rpm at 4°C for 10 minutes and the supernatant was loaded onto 10% to 50% sucrose gradients, followed by ultracentrifugation for 2.5h, at 35,000rpm (200,000*g*) at 4°C with SW41 Ti rotor (Beckman).4C$^0$ . Absorbance at 260nm was recorded using a Piston Gradient Fractionator equipped with a TRIAX full spectrum flow cell (BioComp). Polysome to monosome (P/M) ratios were calculated by comparing the areas under the monosome and polysome peaks. Quantification of various ribosomal fractions: 40S, 60S, 80S, heavy polysomes, according to wt distribution[15,16].

***Gut integrity:*** Young female flies from the indicated genotype were collected into a fresh vial (10 flies per vial), and kept in a humidified, temperature-controlled incubator at 29°C for seven days. Gut integrity was determined using the "smurf assay" : Flies were fed with blue-colored food and its spread to the entire abdomen was determined [6].

***Overall fly survival.*** Flies were reared at 29C$^0$ and transferred into vials containing fresh food every two days. Viability was measured at the indicated time points and LT50 was determined (time in days at which 50% of the fly population died). Statistical analysis was calculated using the GraphPad Prism 5.00 (GraphPad Software, San Diego, CA, USA).

**Statistical analysis:** In all experiments data were collected from at least three independent experiments. Statistical analysis, z-test comparisons were performed using Prism6 ANOVAs software. Significance is indicated by **** = P < 0.0001, ***P < 0.001; **P <0.01; *=P <0.1.

Fig S1 - Daniel *et.al*

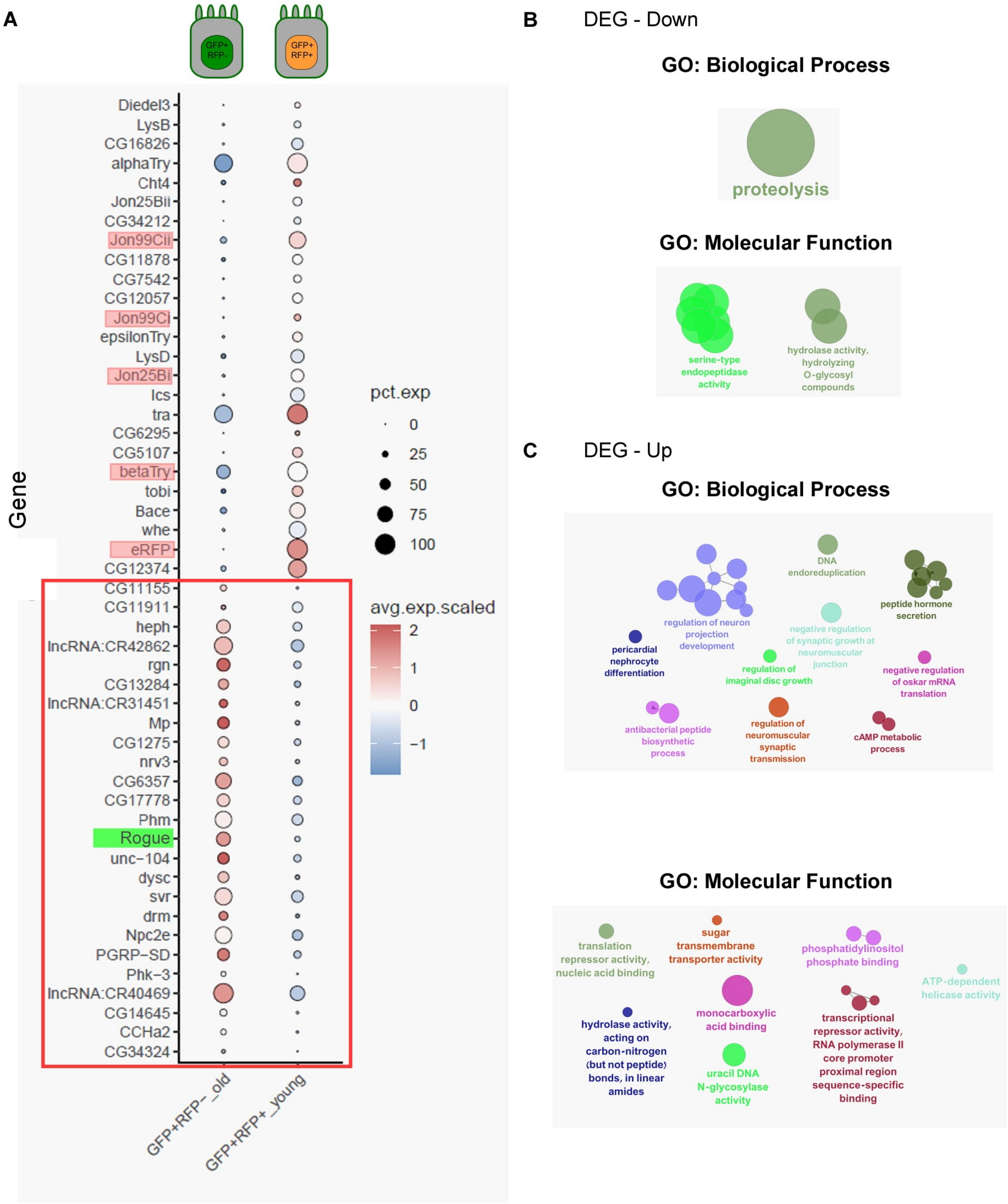

Fig S2 - Daniel *et.al*

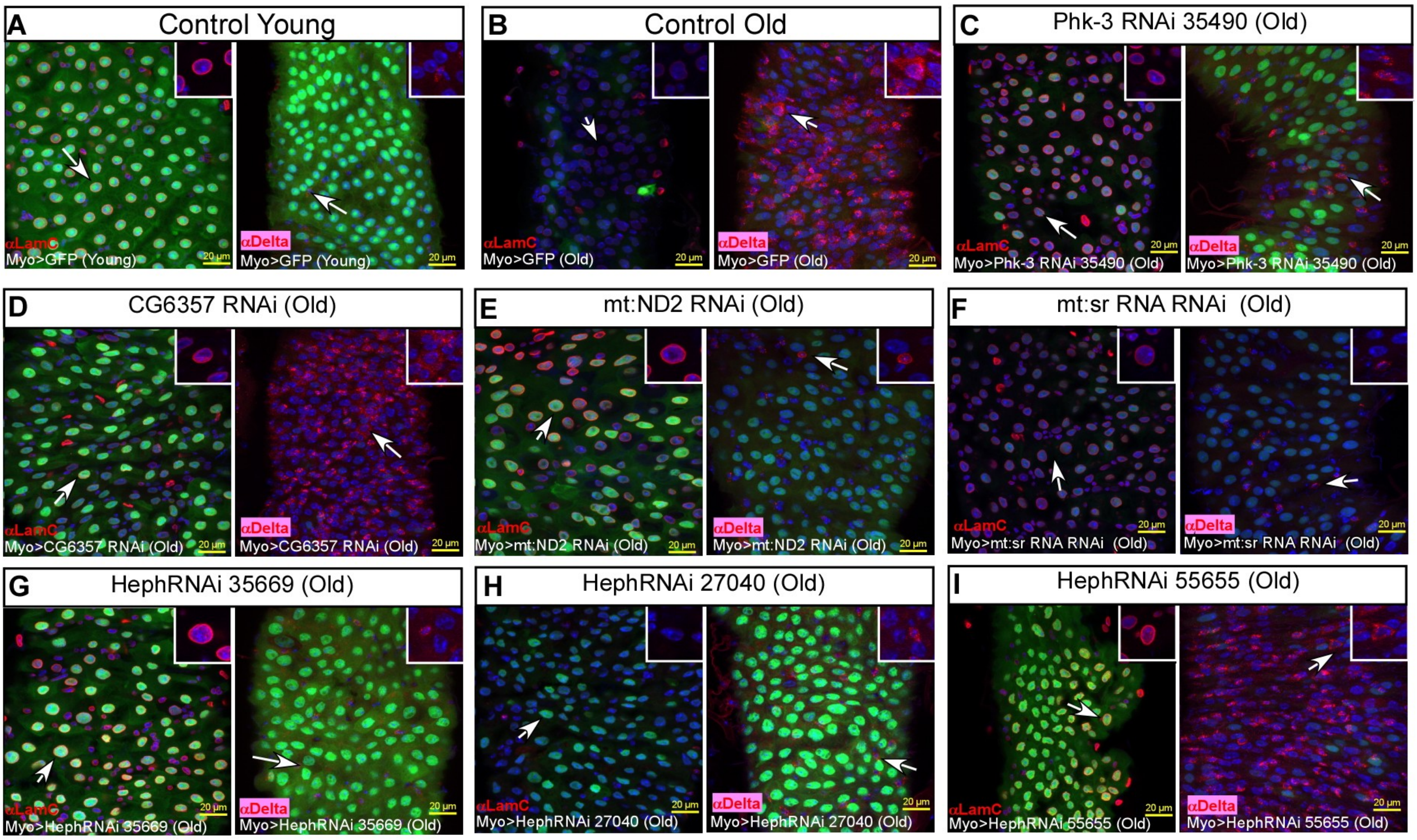


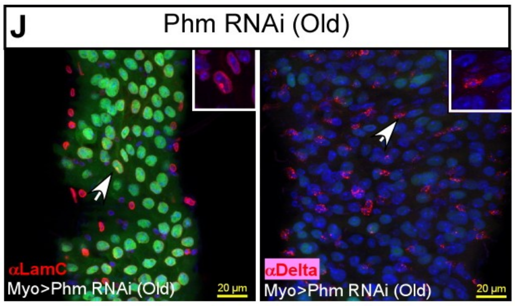


K

| Gene RNAi in Old | LamC exp. | Delta exp. | Gene RNAi in Old | LamC exp. | Delta exp. |
|---|---|---|---|---|---|
| phk-3 | full rescue | partial rescue | phm | minimal rescue | partial rescue |
| mt:ND2 | partial rescue | partial rescue | Heph 55655 | partial rescue | no rescue |
| mt:srRNA | full rescue | full rescue | Heph 35669 | full rescue | partial rescue |
| CG6357 | minimal rescue | no rescue | Heph 27040 | no rescue | minimal rescue |
| mt:ND3 | full rescue | full rescue | Svr | full rescue | full rescue |

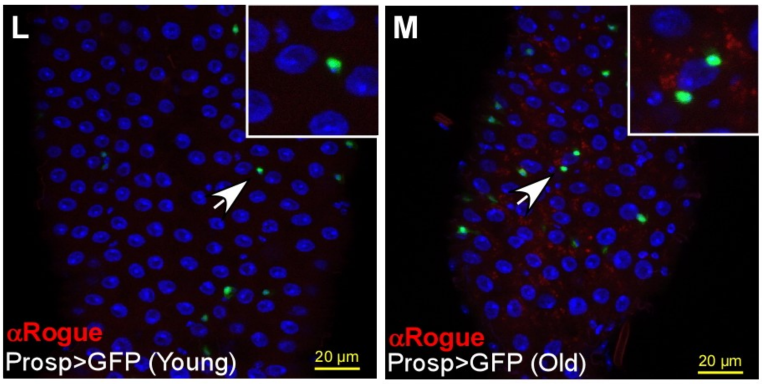

Fig. S3 - Daniel *et.al*

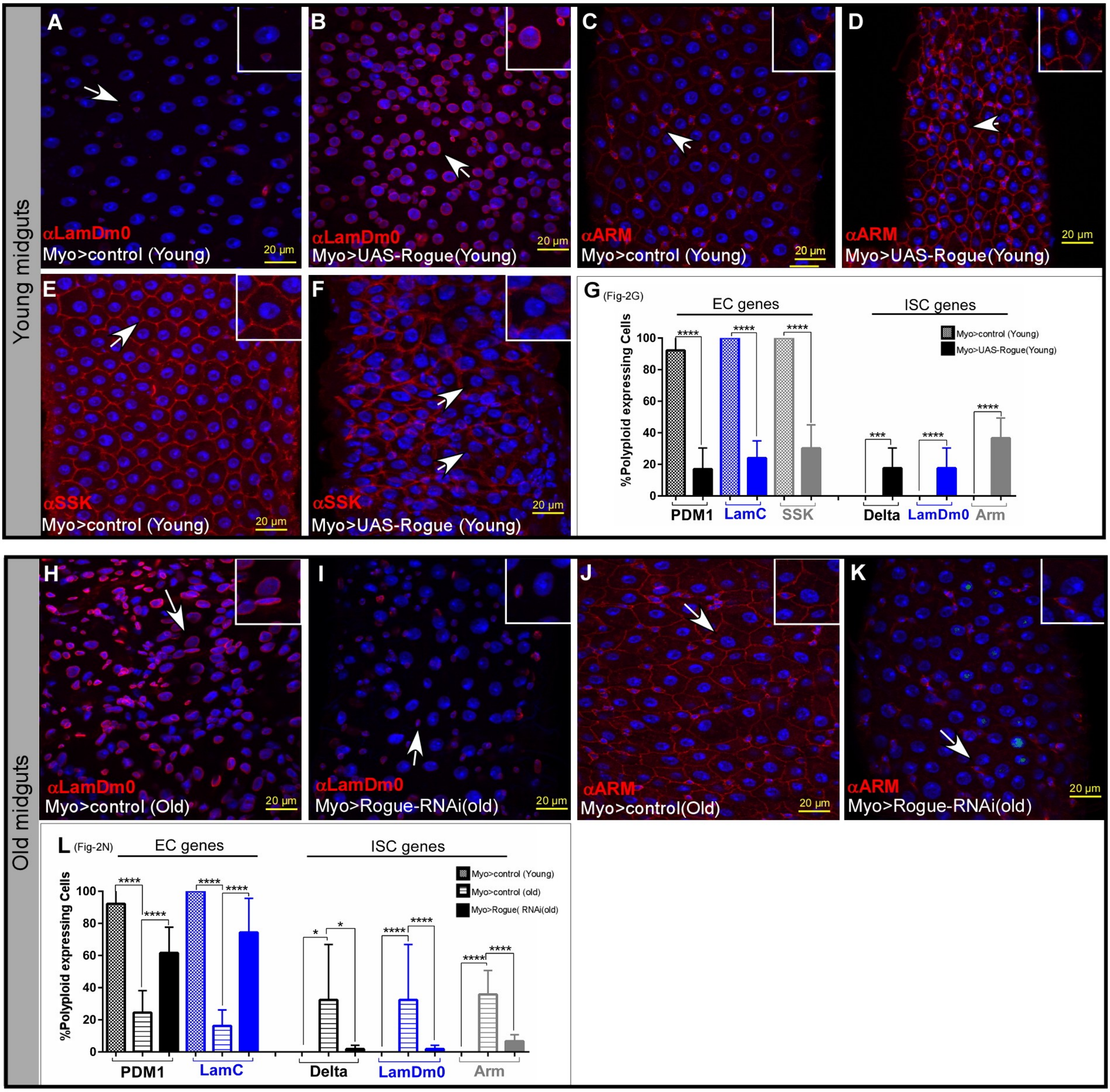

Fig. S4- Daniel *et.al.*

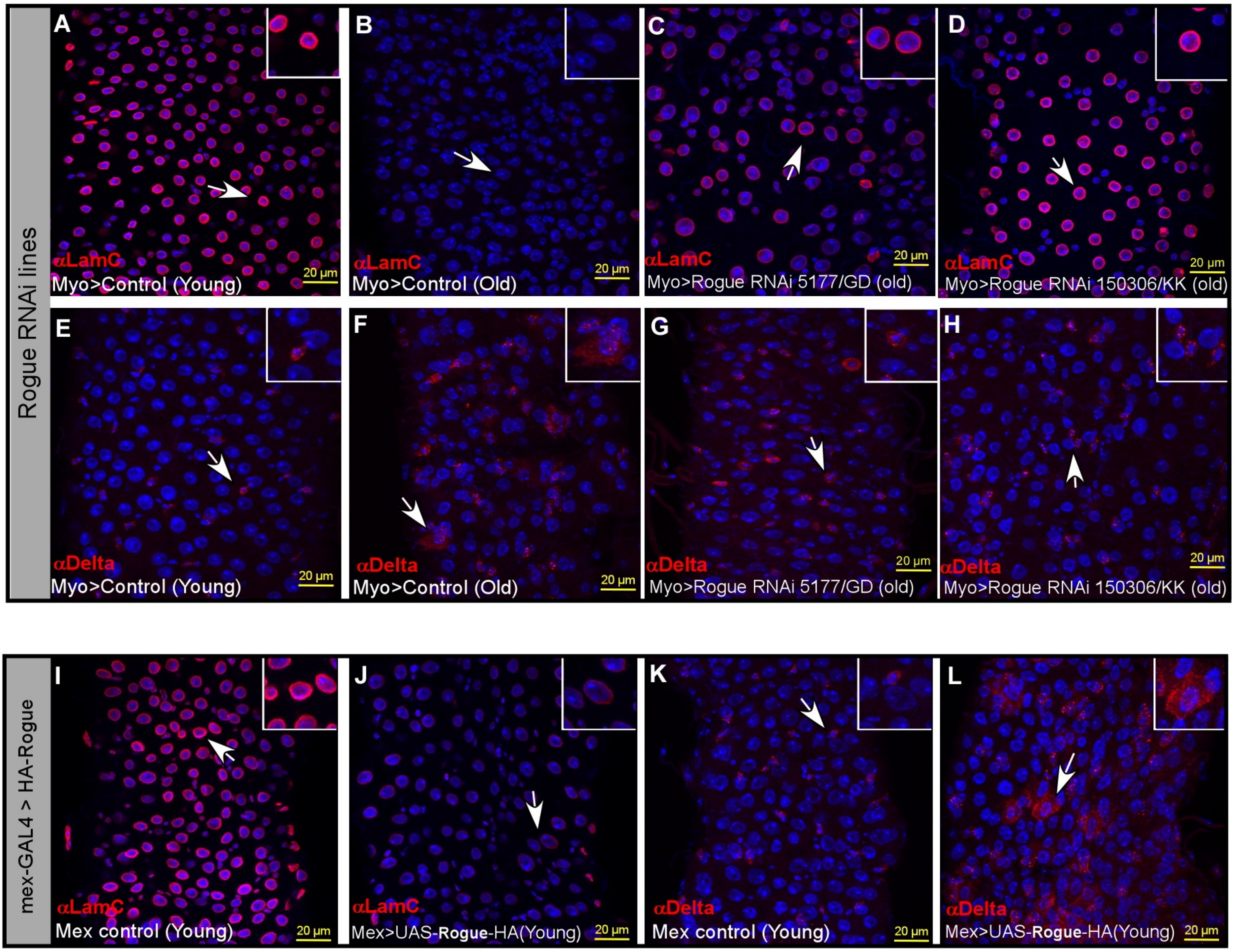

Fig. S5: Daniel *et.al.*

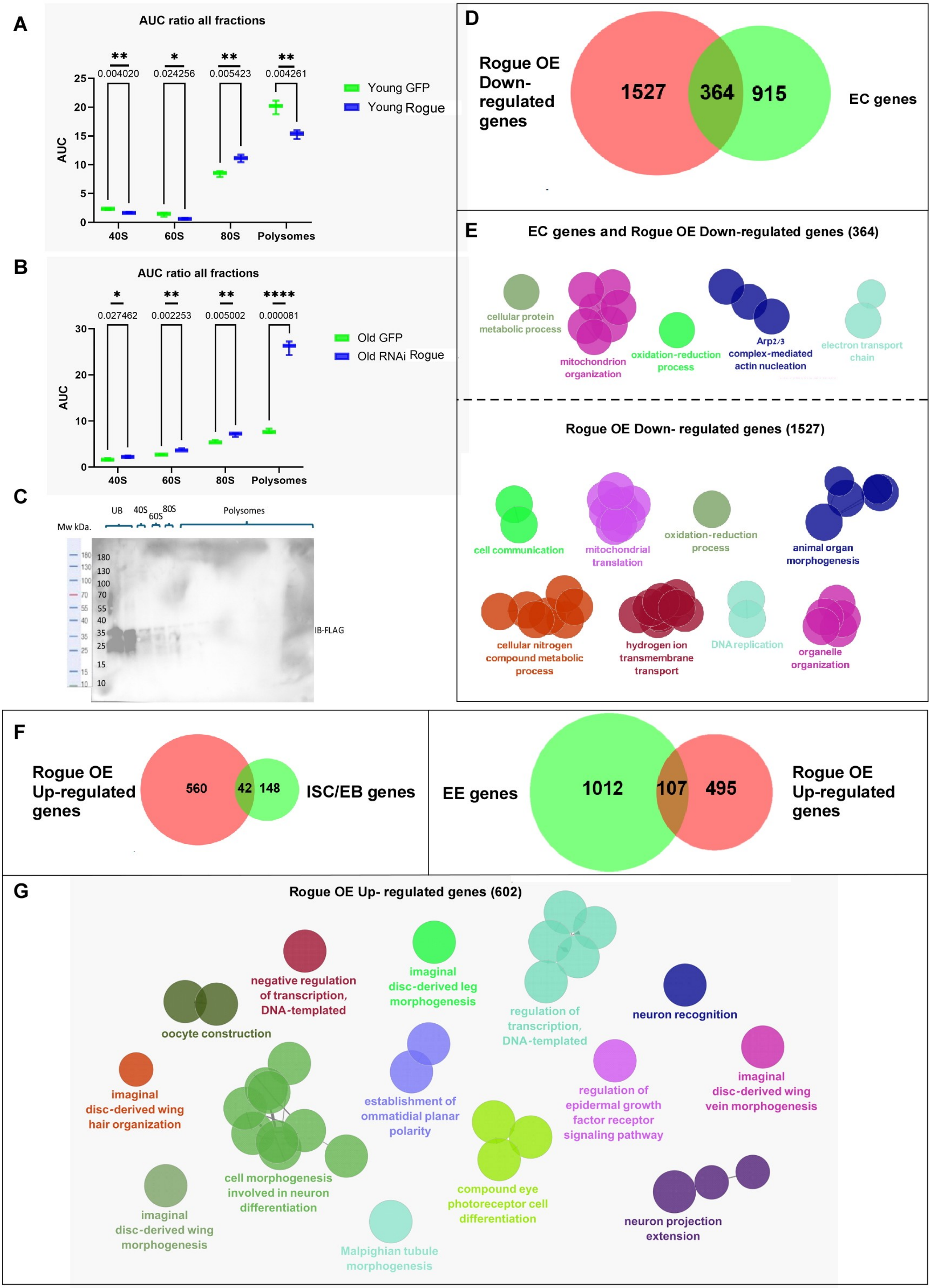

Fig. S6 - Daniel *et.al.*

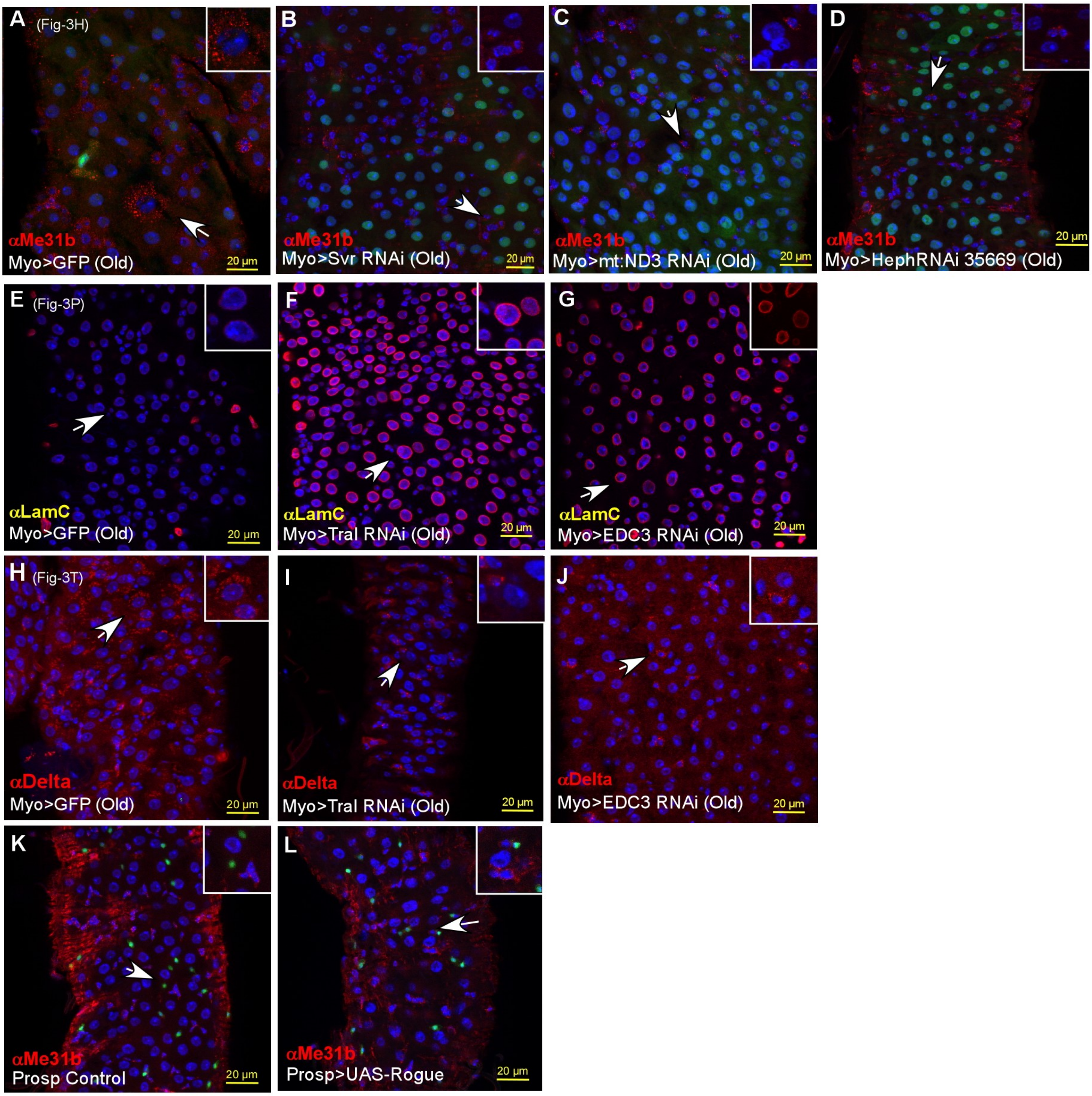

Fig. S7- Daniel *et.al.*

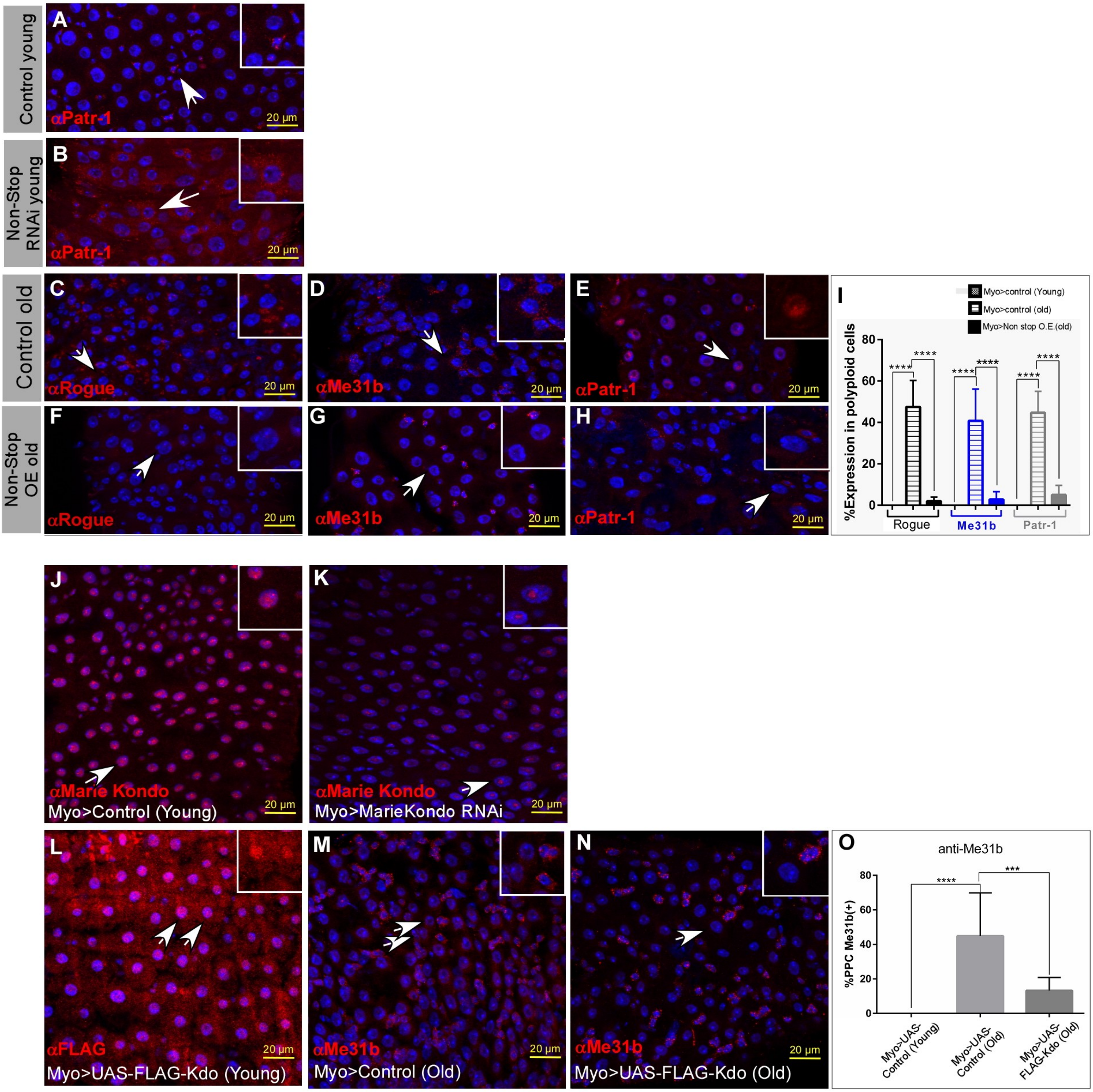

Fig. S8 - Daniel *et.al.*

**A**

| Gene | CG# | FBgn | DIFFERENTIATION | Single cell |
|---|---|---|---|---|
| | | | | (GFP+RFP-old) Vs (RFP+GFP+ young) |
| Lamin Dm0 | CG6944 | FBgn0002525 | Down | #N/A |
| Lamin C | CG10119 | FBgn0010397 | #N/A | #N/A |
| Otefin | CG5581 | FBgn0266420 | Down | #N/A |
| Odd skiped | CG3851 | FBgn0002985 | UP | #N/A |
| nubbin (Pdm) | CG34395 | FBgn0085424 | UP | #N/A |
| Delta | CG3619 | FBgn0000463 | #N/A | #N/A |
| Recq4 | CG7487 | FBgn0040290 | Down | #N/A |
| γ-Tubulin at 37C | CG17566 | FBgn0010097 | #N/A | #N/A |
| γ-Tubulin at 23C | CG3157 | FBgn0260639 | Down | #N/A |
| maelstorm | CG11254 | FBgn0016034 | Down | #N/A |
| Rogue | CG13928 | FBgn0035246 | Up | Up |
| UbcE2H | CG2257 | FBgn0029996 | #N/A | #N/A |
| kaz | CG31357 | FBgn0051357 | #N/A | #N/A |
| soujy | CG3295 | FBgn0034573 | #N/A | #N/A |
| ranbpm | CG42236 | FBgn0262114 | #N/A | #N/A |
| muskelin | CG8811 | FBgn0033757 | #N/A | #N/A |
| cg7611 | CG7611 | FBgn0037094 | #N/A | #N/A |

| Gene | CG# | FBgn | DIFFERENTIATION | Single cell |
|---|---|---|---|---|
| | | | | (GFP+RFP-old) Vs (RFP+GFP+ young) |
| me31b | CG4916 | FBgn0004419 | #N/A | #N/A |
| Houki | CG6617 | FBgn0030944 | #N/A | #N/A |
| esg | CG3758 | FBgn0001981 | Down | Up |
| Pros | CG17228 | FBgn0004595 | #N/A | #N/A |
| betatry | CG18211 | FBgn0010357 | #N/A | Down |
| lambdatry | CG12350 | FBgn0043470 | Up | #N/A |
| Patr-1 | CG5208 | FBgn0266053 | #N/A | #N/A |
| silver | CG4122 | FBgn0004648 | Up | Up |
| Pherokine 3 | CG9358 | FBgn0035089 | #N/A | Up |
| CG6357 | CG6357 | FBgn0033875 | Down | Up |
| Ribosomal protein S14a | CG1524 | FBgn0004403 | #N/A | Up |
| charybde | CG7533 | FBgn0036165 | #N/A | #N/A |
| trailer hitch | CG10686 | FBgn0041775 | #N/A | #N/A |
| Edc3 | CG6311 | FBgn0036735 | #N/A | #N/A |
| Ge-1 | CG6181 | FBgn0283682 | Down | #N/A |
| yipee | CG1989 | FBgn0288857 | Up | #N/A |

### 2. Supp. Fig. S1-7 legends

**Figure S1: Results of sc-RNA-seq combined with G-TRACE. (A)** Examples of EC genes exhibiting reduced or increased expression in aged PPC**$^{RFP-, GFP+}$ compared to young ECs$^{RFP+GFP+}$ cells (•, • mark high and low expression respectively). Genes marked in pink indcate genes that are bona fide ECs genes. Red box denotes examples of genes exhibiting increase expression in PPC**$^{GFP+, RFP-}$ compared to EC $^{GFP+, RFP+}$. Green box highlights the translational repressor Rogue (CG13928). **(B, C)** Cytoscape analysis of molecular function and biological process downregulated (B), or upregulated (C) in PPC**$^{RFP-, GFP+}$ compared to young ECs$^{RFP+GFP+}$.

**Figure S2: Impact of eliminating potential IBs from aged ECs on EC identity . (A-K)** RNAi-mediated elimination of genes exhibiting ectopic mRNA expression in PPC** (FigS1A red box), greatly restores cell identity of aged EC-like cells. Confocal images of young midguts (A) and five weeks old-aged midgut tissue expressing control (B; adopted from Fig.1I, J), or the indicated UAS-RNAi transgenes (C-J) expressed in aged ECs using the MyoIA::Gal4/Gal80ts. MyoIA>UAS-GPF marks fully differentiated ECs, LamC or Delta are shown in red as indicated, and DAPI (blue) marks DNA. White arrows point to cells shown in insets and scale bar is 20μM. **(K)** Qualitative summary of the extent of protein level restoration for each gene based on multiple biological repeats (n>3). **(L, M)** Rogue (CG13928, red) is not expressed in young or old EEs. EEs are marked using *prospero*::GAL4>UAS-GFP. Arrows points to cells shown in insets, DAPI marks DNA, scale bar is 20μM

**Figure S3: Rogue is a bona-fide IB. (A-G)** Expression of Rogue in young ECs unlocks EC identity**.** Confocal images of young midguts expressing control or UAS-FLAG-Rogue in ECs using the MyoIA::GAL4ts80 system and the indicated antibodies. **(H-L)** Conditional elimination of Rogue from five weeks old, aged ECs using UAS-RNAi and MyoIA::GAL4ts80 suppresses the ectopic expression of the indicated ISC-related genes. Arrows points to cells shown in insets, and DAPI (blue) marks DNA. (A, B) LamDm0 (C,D ) Arm, (E, F) SSK, (H, I) LamDm0, (J, K) Arm. Quantification of three independent biological repeats are shown G, and L and are adopted form Fig2.

**Figure S4: Validations using additional UAS-Rogue RNAi, HA-UAS-Rogue-transgenic lines and the EC-specific Mex>Gal4 line. (A-H)** Confocal images of young (A) and old midguts (B-H) expressing the indicated UAS- Rogue RNAi transgenes and the indicated antibodies. (C, G) and (D, H) are different UAS-RNAi transgenic lines towards Rogue. **(I-L)** Confocal images of young EC expressing control or UAS-Rogue-HA transgene under the control of Mex>Gal4 that drives expression solely in ECs in the entire adult fly resulted in decline in LamC (I, J) and ectopic Delta expression (K, L).

**Figure S5: Rogue impact ribosomal profiling and its expression cancels EC signature while promoting ISC, EE, and other cell fate expression signatures. Rogue** affects polysome/monosome rations and is associated with 40s, 60s ribosomal fractions. **(A, B)** Quantification of ribosomal fractions: 40S, 60S, 80S, and heavy polysomes in extracts derived from young flies expressing control or UAS-FLAG-Rogue in ECs (A), or old flies expressing control or UAS-Rogue RNAi (B). Box plots are derived from of area under the curves of the corresponding main figures Fig. 3B, 3D. **(C)** Western blot analysis of sucrose gradient fractions of midguts derived monosomes and polysomes expressing UAS-FLAG-Rogue. Rogue is present in the unbound and 40S and 60S fractions. **(D-G)** GTRACE coupled scRNAseq comparisons of

Rogue overexpression in ECs with gene signatures of EC, ISC and EEs, based on: https://www.flyrnai.org/tools/rna_seq_base/web/showProject/query_marker_report/23/plot_coord=1/sample_id=all . (D) Venn Diagram comparison of genes down regulated upon expression of Rogue in ECs to EC gene signature. (E) Upper panel: Cytoscape analysis of molecular and cellular processes down regulated genes shared between Rogue and ECs. Lower panel: Cytoscape analysis of molecular and cellular processes unique to the expression of Rogue and not ECs . (F) Venn Diagram comparisons of genes up regulated upon expression of Rogue in ISC/EBs (left panel) or EEs (right panel). (G) Cytoscape analysis of molecular and cellular processes that are transcriptionally induced by over-expression of Rogue in ECs.

**Figure S6: IBs identify in aging ECs induce p-body proteins that are potential IBs.**
(**A-D)** Examples where expression of the indicated UAS-IB~RNAi in EC using the MyoIA-GAL4ts80 system in five weeks old ECs cancels the ectopic expression of Me31B observed in aged ECs (A). **(E-J)** p-body proteins are potential IBs. Confocal images of midguts where the indicated p-body proteins were eliminated using UAS-RNAi from EC and the MyoIA-GAL4ts80 system in five weeks old ECs restores LamC protein expression and suppress the ectopic expression of Delta protein . Arrows points to cells shown in insets. **(K, L)** Expression of Rogue in EEs does not result in expression of Me31B in Ees or ECs. Arrows point to cells in the insets and quantification is shown in J and N, n=3 ****=p<0.0001.

**Figure S7: Non-stop and Kdo/CTLH E2/E3 complex negatively regulate Pb-proteins. (A, B)** Elimination of Non-stop (Not) from young ECs results in ectopic expression of Patr-1 protein in ECs. Quantification is shown in Fig. 4E. **(C-I)** Expression of Non-stop, but not control, in aged ECs suppresses the age-associated ectopic expression in ECs of Rogue (C, F), Me31b (D, G) or

Patr-1 (E, H) and quantification is shown in (I). **(J, K)** Endogenous Kdo protein is highly expressed in young ECs (J), and its level declines using UAS-Kdo RNAi (K). **(L-O)** Expression of UAS-FLAG-Kdo transgene (red) using the MyoIA-GAL4ts80 system in young EC (L). (M-O) UAS-FLAG Kdo expression, but not control, suppresses the ectopic expression of Me31b observed in aged EC-like cells, arrows point to cells shown in insets, n=3, ***=P<0.0001.

**Figure S8: Regulation of the differentiated state of ECs by CTLH and Rogue.** Table depicting the changes in mRNA expression of the indicated genes upon differentiation of progenitor cells to ECs [6].

**3. Table S1.** Excell file of differentially expressed genes (DEGs) comparing $PPC^{GFP+RFP-}$ young $EC^{\mathbf{GFP+RFP+}}$ or EC over-expressing Rouge using GTRACE-coupled scRNA-seq.

### 4. Resource table:

| Reagent type (species) or resource | Designation | Source or reference | Identifiers | Additional information |
|---|---|---|---|---|
| Genetic reagent (D. melanogaster) | w; *Myo1A*- Gal4; tub -Gal80ts, UAS-GFP | Edgar Bruce lab | | |
| Genetic reagent (D. melanogaster) | w; Prospero-Gal4 | Edgar Bruce lab | | |
| Genetic reagent (D. melanogaster) | w; *Dl*- Gal4/TM6, Tb | Edgar Bruce lab | | |
| | Mex-GAL4 | Claire M. Thomas | | |
| Genetic reagent (D. melanogaster) | UAS-Non-stop RNAi | VDRC | 45775/GD ; 45776/GD | |
| Genetic reagent (D. melanogaster) | UAS-GFP | Bloomington | # 1521 | |
| Genetic reagent (D. melanogaster) | UAS-Non-stop (3rd) | Ryan D Mohan | 2347 | |
| Genetic reagent (D. melanogaster) | "G-TRACE" (w*; P{UAS-RedStinger}6, P{UAS-FLP.Exel}3, P{Ubi-p63E(FRT.STOP)Stinger}15F2.) | Bloomington | #28281 | |
| Genetic reagent (D. melanogaster) | pUAST - FLAG-CG13928 | Kausik Si | | |
| Genetic reagent (D. melanogaster) | pUAST - FLAG-CG13928 | Kausik Si | | |
| Genetic reagent (D. melanogaster) | UAS-CG13928 RNAi | Kausik Si | #28646 | |
| Genetic reagent (D. melanogaster) | pUAST-Orb2B-HA | Kausik Si | | |
| Genetic reagent (D. melanogaster) | UAS-EDC3-RNAi | Bloomington | 28584 | |
| Genetic reagent (D. melanogaster) | UAS-srl RNAi (FBgn0037248) | Bloomington | 33914 | |
| Genetic reagent (D. melanogaster) | UAS-Patr-1 (FBgn0266053) /TM3 RNAi | Bloomington | 34667 | |
| Genetic reagent (D. melanogaster) | UAS-tral RNAi (FBgn0041775) | Bloomington | 38908 | |
| Genetic reagent (D. melanogaster) | UAS-me31B RNAi (FBgn0004419) | Bloomington | 28566 | |
| Genetic reagent (D. melanogaster) | UAS-CG13928 RNAi | Bloomington | 28646 | |
| Genetic reagent (D. melanogaster) | CG4612-RNAi | Bloomington | 52497 | |
| Genetic reagent (D. melanogaster) | UAS-orb2 RNAi (FBgn0264307) | Bloomington | 60424 | |
| Genetic reagent (D. melanogaster) | UAS-orb2 RNAi (FBgn0264307) | Bloomington | 27050 | |
| Genetic reagent (D. melanogaster) | UAS-Ranbpm RNAi | Bloomington | 61172 | |
| Genetic reagent (D. melanogaster) | CG3295 RNAi (FBgn0034573) - UAS-Souji RNAi | Bloomington | 61896 | |
| Genetic reagent (D. melanogaster) | UAS-UbcE2H RNAi (FBgn0029996) - UBC-E2H/UAS-Marie Kondo RNAi | Bloomington | 51410 | |
| Genetic reagent (D. melanogaster) | UAS-Kaz RNAi | Bloomington | 40248 | |
| Genetic reagent (D. melanogaster) | UAS-mt:srRNA RNAi | Bloomington | 41583 | |
| Genetic reagent (D. melanogaster) | UAS-heph RNAi (FBgn0011224) | Bloomington | 35669 | |
| Genetic reagent (D. melanogaster) | UAS-Phm RNAi (FBgn0283509) | Bloomington | 55392 | |
| Genetic reagent (D. melanogaster) | UAS-heph RNAi(FBgn0011224) | Bloomington | 27040 | |
| Genetic reagent (D. melanogaster) | UAS-heph RNAi(FBgn0011224) | Bloomington | 55655 | |
| Genetic reagent (D. melanogaster) | UAS-mt:ND3 RNAi(FBgn0013681) | Bloomington | 77153 | |
| Genetic reagent (D. melanogaster) | UAS-mt:ND5 RNAi(FBgn0013684) | Bloomington | 77336 | |
| Genetic reagent (D. melanogaster) | UAS-Phk-3 RNAi | Bloomington | 35490 | |
| Genetic reagent (D. melanogaster) | UAS-CG6357 RNAi(FBgn0033875) | Bloomington | 43151 | |
| Genetic reagent (D. melanogaster) | UAS-mt:ND2 RNAi(FBgn0013680) | Bloomington | 77154 | |
| Genetic reagent (D. melanogaster) | w[1118]; P{w[+mC]=UAS-Prosbeta6[1].B}2B; P{UAS-Prosbeta2[1]}1B | Steller | 417, DTS5; DTS7 | |
| **Reagent type (species) or resource** | **Designation** | **Source or reference** | **Identifiers** | **Additional information** |
| Genetic reagent (D. melanogaster) | UAS-svr RNAi (FBgn0004648) | Bloomington | 44487 | |
| Genetic reagent (D. melanogaster) | UAS-svr-GAL4 | Bloomington | 31413 | |
| Genetic reagent (D. melanogaster) | UAS-CG13928-HA | Fly ORF | F002460 | |
| Genetic reagent (D. melanogaster) | UAS-CG13928 RNAi | VDRC | 105306/KK | |
| Genetic reagent (D. melanogaster) | UAS-CG13928 RNAi | VDRC | 5177/GD | |
| Genetic reagent (D. melanogaster) | UbcE2H-HAFF (1) | Ryan | C-TERM CG2257 | |
| Genetic reagent (D. melanogaster) | UbcE2H-HAFF/Tm3 (4) | Ryan | N-TERM CG2257 | |
| Antibody | anti-Armadillo (Mouse monoclonal) | DHSB | N2 7A1 Armadillo | (1:500) |
| Antibody | anti-Delta (Mouse monoclonal(IgG1)) | DHSB | C594.9B | (1:50) |
| Antibody | anti-MESH(Rabbit polyclonal) | Mikio Furuse lab | | (1:100) |
| Antibody | anti-SSK (Rabbit polyclonal) | Mikio Furuse lab | | (1:100) |
| Antibody | anti Lamin C (Mouse monoclonal) | Yossef Gruenbaum lab | | (1:500) |
| Antibody | anti Otefin (Mouse monoclonal) | Yossef Gruenbaum lab | | (1:10) |
| Antibody | anti-Lamin Dm0(Rabbit polyclonal) | Yossef Gruenbaum lab | | (1:300) |
| Antibody | Guinee pig anti-Coilin | Joseph Gall lab | | (1:2000) |
| Antibody | anti-Non-stop (Rabbit polyclonal) | Cloud et al. 2019 | | (1:100) |
| Antibody | anti-Pdm1 (Rabbit polyclonal) | Di'az-Benjumea lab | | (1:50) |
| Antibody | anti-CG13928 (Guinee pig) | Kausik Si | | (1:50) |
| Antibody | anti-Me31b (Mouse monoclonal) | Nakamura, Akira | | (1:500) |
| Antibody | anti-Patr-1 (Rabbit polyclonal) | Nakamura, Akira | | (1:500) |
| Antibody | anti-orb (Guinee pig) | Kausik Si | | (1:100) |
| Antibody | anti-RanBPM (Rabbit polyclonal) | Olivia Rissland | | (1:500) |
| Antibody | anti-Houki (Rabbit polyclonal) | Olivia Rissland | | (1:500) |
| Antibody | anti-Muskelin (Rabbit polyclonal) | Olivia Rissland | | (1:500) |
| Antibody | anti-Kaz (Rabbit polyclonal) | Olivia Rissland | | (1:500) |
| Antibody | anti-flag (Mouse monoclonal) | sigma | | (1:1000) |
| Antibody | Alexa Fluor® 568 goat anti-mouse IgG1(γ1 | invitrogen | A21124 | (1:1000) |
| Antibody | Alexa Fluor® 568 goat anti-mouse IgG (H+L) | invitrogen | A11031 | (1:1000) |
| Antibody | Alexa Fluor® 568 goat anti-rabbit IgG (H+L) | invitrogen | A11036 | (1:1000) |

### 5 References:


1. Bischof, J., Maeda, R.K., Hediger, M., Karch, F., and Basler, K. (2007). An optimized transgenesis system for Drosophila using germ-line-specific φC31 integrases. Proceedings of the National Academy of Sciences of the United States of America *104*, 3312–3317. https://doi.org/10.1073/PNAS.0611511104.

2. Brand, A.H., and Perrimon, N. (1993). Targeted gene expression as a means of altering cell fates and generating dominant phenotypes. Development *118*, 401–415. https://doi.org/10.1242/dev.118.2.401.

3. Salmeron, J.M., Leuther, K.K., and Johnston, S.A. (1990). GAL4 mutations that separate the transcriptional activation and GAL80-interactive functions of the yeast GAL4 protein. Genetics *125*, 21–27. https://doi.org/10.1093/genetics/125.1.21.

4. Jiang, H., Patel, P.H., Kohlmaier, A., Grenley, M.O., McEwen, D.G., and Edgar, B.A. (2009). Cytokine/Jak/Stat Signaling Mediates Regeneration and Homeostasis in the Drosophila Midgut. Cell *137*, 1343–1355. https://doi.org/10.1016/j.cell.2009.05.014.

5. Erez, N., Israitel, L., Bitman-Lotan, E., Wong, W.H., Raz, G., Cornelio-Parra, D.V., Danial, S., Flint Brodsly, N., Belova, E., Maksimenko, O., et al. (2021). A Non-stop identity complex (NIC) supervises enterocyte identity and protects from premature aging. eLife *10*, e62312. https://doi.org/10.7554/eLife.62312.

6. Flint Brodsly, N., Bitman-Lotan, E., Boico, O., Shafat, A., Monastirioti, M., Gessler, M., Delidakis, C., Rincon-Arano, H., and Orian, A. (2019). The transcription factor Hey and nuclear lamins specify and maintain cell identity. eLife *8*, e44745. https://doi.org/10.7554/eLife.44745.

7. Evans, C.J., Olson, J.M., Ngo, K.T., Kim, E., Lee, N.E., Kuoy, E., Patananan, A.N., Sitz, D., Tran, P., Do, M.-T., et al. (2009). G-TRACE: rapid Gal4-based cell lineage analysis in Drosophila. Nat Methods *6*, 603–605. https://doi.org/10.1038/nmeth.1356.

8. Shaw, R.L., Kohlmaier, A., Polesello, C., Veelken, C., Edgar, B.A., and Tapon, N. (2010). The Hippo pathway regulates intestinal stem cell proliferation during *Drosophila* adult midgut regeneration. Development *137*, 4147–4158. https://doi.org/10.1242/dev.052506.

9. Hung, R.J., Li, J.S.S., Liu, Y., and Perrimon, N. (2021). Defining cell types and lineage in the Drosophila midgut using single cell transcriptomics. Current opinion in insect science *47*, 12–17. https://doi.org/10.1016/J.COIS.2021.02.008.

10. ggplot2: Elegant Graphics for Data Analysis (3e) https://ggplot2-book.org/.

11. Pedersen T (2022). patchwork: The Composer of Plots.

12. Zheng, G.X.Y., Terry, J.M., Belgrader, P., Ryvkin, P., Bent, Z.W., Wilson, R., Ziraldo, S.B., Wheeler, T.D., Mcdermott, G.P., Zhu, J., et al. (2017). Massively parallel digital transcriptional profiling of single cells. nature.com. https://doi.org/10.1038/ncomms14049.

13. Satija, R., Farrell, J.A., Gennert, D., Schier, A.F., and Regev, A. (2015). Spatial reconstruction of single-cell gene expression data. Nat Biotechnol *33*, 495–502. https://doi.org/10.1038/nbt.3192.

14. Stuart, T., Butler, A., Hoffman, P., Hafemeister, C., Papalexi, E., Mauck, W.M., Hao, Y., Stoeckius, M., Smibert, P., and Satija, R. (2019). Comprehensive Integration of Single-Cell Data. Cell *177*, 1888-1902.e21. https://doi.org/10.1016/j.cell.2019.05.031.

15. Ngoc, H., Luong, B., Kalogeridi, M., Vontas, | John, and Denecke, S. (2022). Using tissue specific P450 expression in Drosophila melanogaster larvae to understand the spatial distribution of pesticide metabolism in feeding assays. Wiley Online LibraryHNB Luong, M Kalogeridi, J Vontas, S DeneckeInsect Molecular Biology, 2022•Wiley Online Library ,*31* 376–369. https://doi.org/10.1111/imb.12765.

16. Shiber, A., Döring, K., Friedrich, U., Klann, K., Merker, D., Zedan, M., Tippmann, F., Kramer, G., and Bukau, B. (2018). Cotranslational assembly of protein complexes in eukaryotes revealed by ribosome profiling. Nature *561*, 268–272. https://doi.org/10.1038/s41586-018-0462-y.
